# Inhibitory neurons marked by the connectivity molecule Kirrel3 regulate memory precision

**DOI:** 10.1101/2024.07.05.602304

**Authors:** Arnulfo Tuñon-Ortiz, Dimitri Tränkner, Chandler M. Peterson, Omar Shennib, Fangfei Ye, Jiani Shi, Sarah N. Brockway, Olivia Raines, Abbey Mahnke, Matthew Grega, Keun-Young Kim, Mark H. Ellisman, James G. Heys, Moriel Zelikowsky, Megan E. Williams

## Abstract

The homophilic adhesion molecule Kirrel3 drives synapse formation between dentate granule (DG) neurons and GABA neurons, and Kirrel3 gene variants are associated with neurodevelopmental disorders in humans. However, the circuit function and behavioral relevance of Kirrel3-expressing neurons are unknown. Using intersectional genetics, we identified a population of Kirrel3-expressing GABA neurons that regulate memory discrimination in male and female mice. Using chemogenetics with in vivo electrophysiology and behavioral assays, we discovered that activating Kirrel3-expressing GABA neurons, but not parvalbumin neurons, potently inhibits CA3 neuron activity and impairs contextual memory discrimination during recall, revealing a critical role for these neurons in the retrieval of precise memories. Light and electron microscopy of Kirrel3-expressing GABA neurons suggests that they receive direct excitation from DG neurons and project onto CA3 dendrites. Together, this multi-scale approach demonstrates how cell type-specific expression of adhesion molecules mark subsets of neurons that control key features guiding memory and behavior.

**SIGNIFICANCE STATEMENT:** Here, we identified a never-before studied group of inhibitory neurons defined by expression of the gene Kirrel3, which is a synaptic cell adhesion molecule and risk gene for neurodevelopmental disorders. We show that Kirrel3 inhibitory neuron activity exerts exceptionally strong control over CA3 neuron activity and impairs the ability of mice to distinguish between different contexts during memory retrieval. Thus, Kirrel3 inhibitory neurons regulate memory precision in mice, and our study employs a powerful framework linking molecular identity and synaptic specificity to behavioral function in the adult brain.

## INTRODUCTION

Synaptic cell adhesion molecules (CAMs) regulate synapse formation, maintenance, and function, with new and unexpected roles continuing to emerge (Südhof 2018). Given these critical functions, it is not surprising that genetic mutations in CAMs are linked to many synaptic disorders in humans, including autism spectrum disorders, intellectual disability, and mental illnesses (Südhof 2017, Südhof 2018). Their strong evolutionary conservation suggests that each CAM has a distinct and essential role in building circuits with precision. However, studying CAMs presents several challenges. They are expressed in relatively low copy numbers per cell, are often restricted to specific neuronal and synaptic subtypes, and their functions can be subtle, transient, or compensated, making them difficult to detect. Despite these challenges, understanding how disease-associated CAMs influence neural circuit function remains a critical goal in neuroscience.

To address these challenges, we adopted an alternative approach to studying synaptic CAMs. Rather than focusing solely on the molecular function of an individual CAM protein using knockout mice, we investigated how neurons expressing a cell type specific CAM regulate local circuit activity. We focused on Kirrel3, a single-pass transmembrane protein in the immunoglobulin superfamily, which is repeatedly identified as a genetic risk factor for neurological disorders that frequently includes autism spectrum disorder and intellectual disability (Bhalla, Luo et al. 2008, Kaminsky, Kaul et al. 2011, Ben-David and Shifman 2012, Guerin, Stavropoulos et al. 2012, Michaelson, Shi et al. 2012, Neale, Kou et al. 2012, Talkowski, Rosenfeld et al. 2012, De Rubeis, He et al. 2014, Iossifov, O’Roak et al. 2014, Wang, Guo et al. 2016, Li, Wang et al. 2017, Guo, Duyzend et al. 2019, Leblond, Cliquet et al. 2019, Taylor, Martin et al. 2020). In vitro, Kirrel3 mediates trans-cellular adhesion and synapse formation (Taylor, Martin et al. 2020). In vivo, Kirrel3 is specifically required for synapse formation between dentate gyrus (DG) granule cells and GABAergic neurons in hippocampal area CA3 of mice (Martin, Muralidhar et al. 2015, Martin, Woodruff et al. 2017, Taylor, Martin et al. 2020, Ciaccio, Leonardi et al. 2021, Querzani, Sirchia et al. 2023). Kirrel3 loss-of-function also leads to increased CA3 pyramidal neuron activity during development (Martin, Muralidhar et al. 2015) and some general behavioral abnormalities in adult mice (Choi, Han et al. 2015, Hisaoka, Komori et al. 2018, Volker, Maar et al. 2018). However, it remains unclear how Kirrel3-expressing neurons contribute to circuit function in the adult brain and how they specifically influence hippocampal behavior. To investigate this, we developed a transgenic mouse model that enables selective labeling and manipulation of Kirrel3-expressing neurons. This model allowed us to directly assess the role of Kirrel3-expressing neurons in circuit function and behavior without the confounding effects of abnormal brain development.

In the hippocampus, Kirrel3 is expressed by glutamatergic DG neurons and a subset of GABAergic (GABA) neurons (Martin, Muralidhar et al. 2015). While DG neuron function in learning and memory is well studied, the role of Kirrel3-expressing GABA (K3-GABA) neurons in CA3 is unknown. We combined intersectional genetics with chemogenetics to selectively activate K3-GABA neurons and assessed their role in contextual fear memory. We found that activating K3-GABA neurons during recall decreases memory discrimination between a learned contextual fear context and a novel neutral context. This effect on memory precision was not observed when parvalbumin-expressing (PV) neurons were activated, suggesting a unique role for K3-GABA neurons in memory discrimination. Further anatomical and functional analyses revealed that K3-GABA neurons receive synaptic input from DG neurons and strongly inhibit CA3 neurons. In contrast to traditional approaches that isolate either molecular function or circuit dynamics, our study employs a powerful framework that links molecular identity, synaptic specificity, and behavioral function in the adult brain. This highlights the validity of defining neurons based on synaptic CAMs, as they are likely to share distinct connectivity patterns. Moreover, because Kirrel3 mutations are linked to neurodevelopmental disorders, our study provides a framework for investigating how Kirrel3 neuron dysfunction alters neural circuits and contributes to cognitive and behavioral impairments.

## MATERIALS AND METHODS

### Mouse lines

All mice are maintained in the C57Bl/6 background. For the Kirrel3-Flp line, FlpO-P2A was inserted in frame and immediately upstream of the Kirrel3 start codon so that Kirrel3 protein expression is unperturbed. We inserted FlpO at the start because the Kirrel3 gene undergoes alternative splicing at its C-terminus (Traenkner, Shennib et al. 2023). Transgenic mice were generated at the University of Nebraska School of Medicine Transgenic core using the Easi-CRISPR method of homologous recombination^45^. Briefly, single cell C57Bl/6 mouse zygotes were injected with preassembled Cas9 ribonucleotide complexes containing Cas9 protein, a Kirrel3 crRNA with the sequence AGGA<u>ATG</u>AGACCTTTCC/AGC (where / indicates cut site and the underline indicates the Kirrel3 start codon), and tracrRNA along with a 1,536 bp single strand DNA (ssDNA) repair template containing the FlpO-P2A sequence and ∼100bp homology arms. A founder mouse was fully sequenced to ensure there were no mutations at insertion sites, and the mouse was bred to a wildtype C57Bl/6 mouse to propagate the strain.

Other mice used are: Gad2-IRES-Cre (JAX stock #010802) (Taniguchi, He et al. 2011), RC::FLTG (JAX stock #026932) (Plummer, Evsyukova et al. 2015), and PV-Cre (JAX stock #017320) (Hippenmeyer, Vrieseling et al. 2005), and C57BL/6J wildtype mice (JAX stock #000664). All mice were maintained and conducted per NIH guidelines on the care and use of animals approved by the University of Utah IACUC committee.

### Immunohistochemistry

Mice were transcardially perfused with 4% paraformaldehyde in 1X phosphate-buffered saline (PBS) prepared from 10X stock consisting of 14.2g of Na2HPO4, 80g of NaCl, 2g KCL, 2g KH2PO4 at a pH of 7.4. Brains were then stored in 4% PFA overnight and cut into 50 µm sections around the hippocampal formation. Sections were incubated in blocking solution (PBS with 3% bovine serum albumin (BSA) and 0.2% Triton-X100) for 1 hour. Sections were incubated overnight with gentle shaking in primary antibody solution, which was prepared with the antibody diluted in PBS with 3% BSA. After 3 washes with PBS, sections were incubated with secondary antibody solution for 2 hours at room temperature, followed by 2 washes in 1xPBS and a final wash in 1xPBS with Hoechst prior to mounting on slides.

### Antibodies

Primary antibodies were used as follows: goat anti-GFP 1:5000 (Abcam, ab6673), rabbit anti-cFos 1:500 (Santa Cruz Biotech, sc-253; Synaptic Systems, 226008), rabbit anti-calretinin 1:2000 (Swant, 7699/3), mouse anti-parvalbumin 1:5000 (Swant, PV235), rat anti-somatostatin 1:500 (Chemicon, MAB354), rabbit anti-calbindin d28k 1:2000 (Swant, CB38a), rabbit anti-VIP 1:500 (Immunostar), rabbit anti-PV 1:250 (Swant, PV27a), rabbit anti-RFP 1:1000 (Fisher, NBP267102), goat anti-mCherry (Novus, NBP3-05558). All secondary antibodies were from Jackson ImmunoResearch Laboratories, were made in donkey, and used at 1:1000.

### Stereotaxic surgeries

Adult mice aged 8 weeks or older were intraperitoneally injected with Buprenorphine (0.1mg/kg) an hour prior to the operation. Mice were anesthetized with oxygen and isoflurane and then readied in the stereotaxic surgery area. An injection of lidocaine (2mg/kg) was administered subcutaneously near the intended incision area which is shaved and sterilized. Small holes were drilled in the skull to expose the region of interest for viral intracranial injection. 500 nL of AAV was delivered bilaterally via picospritzer to the desired region with a 0.03 ms pulse at 60 psi. Upon completion, incisions were sealed with VetBond, and mice were injected subcutaneously with Rimadyl (5 mg/kg carprofen). Mice were allowed to recover on a heating pad and observed for signs of distress before returning to their home cage. Follow-ups continued for three days and consisted of weighing mice and administering Rimadyl to ensure a healthy recovery.

For in vivo electrophysiology experiments, 8 weeks old mice were injected with AAV expressing excitatory (hM3D) DREADDs to activate specific inhibitory neuron types: Kirrel3-Flp/Gad2-IRES-Cre heterozygous mice were injected with AAV ConVERGD hSyn.hM3D-mCherry (pHp.eb), and PV-Cre homozygous mice were injected with AAV hSyn-DIO-hM3D-mCherry. During the injection surgery, a small craniotomy was made upon injection site (coordinate AP -1.6 mm, ML +/-2.3 mm from bregma). Virus was injected bilaterally (500 nl each side) using Nanoject III Injector (Drummond), injected at a rate of 10 nl/s. After injection a custom-designed titanium 572 headplate (9.5 × 33 mm²) was fixed upon skull using Metabond (Parkell). To alleviate pain following surgery, mice were injected subcutaneously with Rimadyl (5mg/kg Carprofen).

### Viruses

Adeno-associated viruses (AAVs) were made using standard iodixanol gradient purification. Viral titers for all viruses used were 10^11 to 10^13. Viruses used are as follows: ConVERGD construct hSyn.hM3Dm.Cherry (PHP.eB) and hSyn.hM3D.HA (AAV8) provided by Lindsay Schwarz lab (Hughes, Pittman et al. 2024); hSyn.DIO.hM3D.mCherry (AAV9) (Addgene 44361); INTRSCT construct hSyn.Cre-on/Flp-on.YFP (PHP.eB) (Addgene 55650) (Fenno, Mattis et al. 2014).

### Fluorescent in situ hybridization (FISH)

Fluorescent in situ hybridization chain reaction was performed as described (Trivedi, Choi et al. 2018). Briefly, mouse brains were cryo-sectioned to 30 μm slices, mounted on slides, fixed in 4% paraformaldehyde (PFA), and washed in PBS. Slices were treated with 1 mg/ml proteinase K in TE buffer and equilibrated in SSC buffer. HCR probes were designed and generated by Molecular Instruments for Gad1, Kirrel3, and GFP. After nuclear staining with Hoechst in PBS, coverslips were mounted in Fluoromount-G (Southern Biotech catalog #0100-01) and imaged on a confocal microscope. For dual in situ/immunohistochemistry, samples were incubated in freshly prepared 4% PFA in 1xPBS made with DEPC water for 10 minutes and washed in 1X DEPC treated PBS (pH 7.4) for 5 minutes at room temperature three times. Samples were blocked for 1 hour at room temperature (100 μL/slide applied over the top of sections) inside a humidified chamber and 75 µL primary antibody solution (anti-mCherry 1:1000) was added per slide and covered with an RNAse free coverslip. Samples were incubated for 1 hour at room temperature and then overnight at 4°C in a humidified chamber. The next day, samples were washed in 1X DEPC treated PBST (pH 7.4) for 5 minutes at room temperature for three washes before 2° antibody was added at 1:1000 then incubated for 1 hour at room temperature inside a humidified chamber. Samples were washed in 1X DEPC treated PBST (pH 7.4) for 5 minutes at room temperature three times and coverslipped with Fluoromount-G mounting reagent.

### Novel environment

Mice were removed from their home cage and intraperitoneally injected with saline or 1.0mg/kg DCZ then immediately placed into a 40 cm x 40 cm plastic container with 5 novel objects evenly spaced apart. Mice were allowed to explore freely for 40 minutes prior to perfusion.

### Contextual Fear Conditioning

Mice underwent a 3-day contextual fear conditioning protocol during their light cycle (6am-6pm). Mice were habituated for 3 days prior to conditioning day (Day 0) for 1 hour in squads of 4 to a neutral habituation room. To create distinct and similar contextual environments, chambers and interchangeable inserts from MedAssociates were used in two rooms. Contexts A and B were presented in the same room, and context C was presented in a different room. Each context was assigned a specific set of features (floor, roof, scent, sound, light, and transport vessels) as described in Extended data Figure 3-1. Context B is meant to closely resemble A, thus only 2 of 6 features (floor and transport) were altered (Extended data Figure 3-1). Mice were injected intraperitoneally (i.p.) with saline or 1.0 mg/kg DCZ (Ferrari, Ogbeide-Latario et al. 2022) 10 minutes prior to being placed into behavior boxes (MedAssociates) on all days or only on a specific day as described in the results section. During acquisition, mice received 4 shocks (1 mA, 2-second duration, 1-minute inter-shock interval), delivered after an initial 3-minute baseline period (Zelikowsky, Bissiere et al. 2013). One minute after the final shock, acquisition ended, and mice were transported back to their home cage. 24 hours later, on test day 1, mice were habituated and injected as described earlier. Conditioned mice were exposed to context A for 8 minutes or to the “similar” context B (counter balanced). Mice were returned to their home cage in the vivarium for a minimum of 5 hours before being tested in the alternative context. Twenty-four hours later, this process was repeated for test day 2, with the distinct, “neutral” context C replacing the “similar” context B. To analyze behavior, component files were made to extract the desired data using automated near-infrared video tracking equipment and computer software (VideoFreeze, MedAssociates). For conditioning day 0, the 3-minutes prior to the first shock were extracted to analyze baseline percent component freezing and motion index. The 30 seconds preceding each shock were also extracted to analyze the percent component freezing for fear acquisition curves. For test days 1 and 2, the percent time freezing was calculated for the entire 8-minute session. All mice used for behavior were verified for correct virus expression and targeting. Mice were excluded from all analyses if they did not express virus in the CA3/hilus region or had more infection outside the CA3/hilus region than inside (Extended Data Figure 3-1).

### cFos analysis

Confocal images (Zeiss 980) were taken at 10X and 20X objective as Z-stacks and tiles, exported as OME-TIFF files, and analyzed in FIJI blind to genotype and condition. Images were processed as follows: z-projected for maximum intensity, despeckled, and split channels. ROIs were drawn for the indicated cell body region. Next the cFos channel background was subtracted by a rolling ball radius value of 15 and then it was subject to thresholding. The thresholded value was determined by opening the threshold window, finding the right-side inflection-point of the histogram curve, and multiplying it by 3. For determining the percent of DREADD-mCherry cells expressing cFos, manual cell counts were conducted after thresholding. For determining the percent of principle cells with cFos, we used the automatic analyze particles function with a circularity range of 0-1.0 and size range of 30 μm^2^-infinity. Counts were verified by merging channels and examining merged fluorescence.

### In vivo electrophysiology and behavioral task

Two days after surgery, mice began water restriction (1 mL of water per day) to maintain 85%-90% of their initial weight during training and recording period. Approximately 5 days after surgery, behavior training began. During the task, mice were head-fixed on a cylindrical treadmill (60-cm circumference, 10-cm width) and learned to navigate through a 3-meter virtual environment projected on five screens surrounding the animal. A solenoid valve was used to deliver water (∼6 μL per drop) via a lick spout. Mice were trained ∼1.5 hours per day until they routinely ran more than 180 meters per hour. The number of days required to reach this criterion varied by mouse and ranged from 7 to 14 days. At least 2 weeks after virus injection, a craniotomy was made over the CA3 injection site and sealed with Kwik-Sil (World Precision Instruments). After at least one day of recovery, we recorded from CA3 acutely using Neuropixel 2.0 while the mouse performed the behavioral task (one recording session per day). Before each recording, the probe was briefly dipped into a fluorescent dye (DiD or DiO, ThermoFisher) to mark the probe track. The probe was lowered using a micromanipulator at 1-2 µm/s, reaching the target depth (typically 2200–2500 µm below the brain surface) after approximately 30–40 minutes. After confirming the target location by a brief recording, we waited for another 15 min to minimize drift during data collection. To control for potential confounds related to movement speed, mice ran on a wheel while their locomotion was monitored in a virtual environment. We first administered a control intraperitoneal (i.p.) injection without liquid, and after 8 minutes, recorded for 15 minutes while the mouse ran. We then administered an i.p. injection of either DCZ solution or saline, waited another 8 minutes to allow the drug to take effect, and recorded for an additional 15 minutes during wheel running. At the end of the recording session, the probe was slowly retrieved at 1-2 um/s and the craniotomy was sealed with Kwik-Sil. The mouse then received a subcutaneously injection of Rimadyl (5mg/kg carprofen) for post-procedure analgesia.

The set-up for the Neuropixel 2.0 system was as described (Bowler, Azhar et al. 2025). Neural recordings were processed on a headstage (HS-2010, Imec) where signals were amplified, multiplexed, filtered, and digitized. These signals were then transferred to an acquisition system (Imec and National Instruments) and streamed into SpikeGLX software (https://billkarsh.github.io/SpikeGLX) for recording. Data were sampled at 30 kHz and digitized with a gain of 100. Behavioral signals, collected using BNC-2110 and PXIe-6341 hardware (National Instruments), were simultaneously streamed into SpikeGLX and time-aligned with the neural data.

Recordings were first preprocessed using CatGT (https://github.com/billkarsh/CatGT), which applied an order-12 band-pass Butterworth filter (300–9000 Hz), removed artifacts, performed phase shift correction, and applied common reference averaging. Motion correction and spike sorting were performed using kilosort4 (Pachitariu, Sridhar et al. 2024). Quality control metrics were obtained using spikeInterface (Buccino, Hurwitz et al. 2020). We classified a well-isolated single unit if it met the following criteria: number of waveform troughs <2, number of waveform peaks <2, firing rate >0.1 Hz and < 6Hz, signal-to-noise ratio >2, firing range <20 (firing range computed by taking the difference between the 95^th^ percentile’s firing rate and the 5^th^ percentile’s firing rate computed over 5s). Isolated putative single units were manually inspected and curated using Phy2 (https://github.com/cortex-lab/phy). The following analysis was done using customized Python scripts. To control for potential confounds due to differences in movement speed, we filtered data based on the animal’s running velocity. For Kirrel3 transgenic mice, data were excluded when running speeds were below 5 cm/s or above 50 cm/s. For PV-Cre mice, data were excluded when running speeds were below 5 cm/s or above 26 cm/s. After data collection, mice were perfused using a 4% PFA solution and the brain was collected for histological analysis. Brains were sectioned at 60 µm thickness and stained with Blue Fluorescent Nissl Stain. Images were collected using VS200 Olympus microscope.

### Correlated light and electron microscopy

A female K3-Flp;Gad2-Cre mouse was infected with an AAV expressing Flp/Cre-dependent GFP. 4 weeks later (at 84 days of age), it was anesthetized with an intraperitoneal injection of ketamine/xylazine and transcardially perfused with a brief flush of Ringer’s solution containing heparin and xylocaine, followed by approximately 50 mL of 0.5% glutaraldehyde/4% paraformaldehyde in 0.15 M sodium cacodylate buffer containing 2 mM CaCl_2_ (CB, pH 7.4). The brain was dissected and post-fixed overnight on ice in the same fixative and prepared 100 µm thick coronal brain sections using a vibratome. GFP expressed in Kirrel3+ GABA neurons bearing hippocampus slices were collected and incubated in DRAQ5 (1:1000, Cell Signaling Technology) on ice for an hour. Confocal images of GFP and DRAQ5 signals in CA3 region of the hippocampus were collected on a confocal microscopy (Olympus FluoView1000; Olympus, Tokyo, Japan) with using 488 nm and 633 nm excitation.

Confocal imaged brain slices were prepared for MicroCT and SBEM as previously described (Bushong, Johnson et al. 2015, Agarwala, Kim et al. , Bouin, Wu et al. 2024). Briefly, immediately after confocal imaging, brain slices were fixed in 2.5% glutaraldehyde in 0.15 M cacodylate buffer (CB, pH 7.4) at 4°C an hour. After removing the fixative, brain slices were washed with 0.15 M CB and then placed into 2% OsO4/1.5% potassium ferrocyanide in 0.15 M CB containing 2 mM CaCl_2_ for 1 h at room temperature (RT). After thorough washing in double distilled water (ddH_2_O), slices were placed into 0.05% thiocarbohydrazide for 30 min. Brain slices were again washed and then stained with 2% aqueous OsO4 for 30 min. Brain slices were washed and then placed into 2% aqueous uranyl acetate overnight at 4°C. Brain slices were washed with ddH2O at RT and then stained with 0.05% en bloc lead aspartate for 30 min at 60°C. Brain slices were washed with ddH_2_O and then dehydrated on ice in 50%, 70%, 90%, 100%, 100% ethanol solutions for 10 min at each step. Brain slices were then washed twice with dry acetone and then placed into 50:50 Durcupan ACM:acetone overnight. Brain slices were transferred to 100% Durcupan resin overnight. Brain slices were then flat embedded between glass slides coated with mold-release compound and left in an oven at 60°C for 72 h.

The MicroCT tilt series were collected using a Zeiss Xradia 510 Versa (Zeiss X-Ray Microscopy) operated at 80 kV (87 µA current) with a x20 magnification and 0.7930 µm pixel size. MicroCT volumes were generated from a tilt series of 2401 projections using XMReconstructor (Xradia). SBEMs were accomplished using Gemini SEM (Zeiss, Oberkochen, Germany) equipped with a Gatan 3 View system and a focal nitrogen gas injection setup. This system allowed the application of nitrogen gas precisely over the block-face of ROI during imaging with high vacuum to maximize the SEM image resolution. Images were acquired in 2.5 kV accelerating voltage and 1 µs dwell time; 6 nm XY pixels, 60 nm Z steps; raster size was 20 k × 20 k, and Z dimension was 1001 images. Volumes were collected using 70% nitrogen gas injection to samples under high vacuum. Once volumes were collected, the images were converted to mrc format and cross-correlation was used for rigid image alignment of the slices using the IMOD image processing package (Kremer, Mastronarde et al. 1996). Tracing and analysis were done manually in VAST-Lite software and exported as .OBJ to Blender to generate the final 3D model.

### Experimental design and statistical analysis

Statistics were calculated in GraphPad Prism. Whenever possible, samples were analyzed blind to genotype or treatment. A full description of sample sizes, statistical tests used, degrees of freedom, F and t values, and effect size as R or eta squared of all statistical comparisons is in Extended Data Table 1. Also included in Extended Data Table 1 are RRIDs for key reagents.

## RESULTS

### Intersectional genetics enables specific targeting and activation of K3-GABA neurons

To study Kirrel3-expressing neurons, we generated a mouse line that expresses Flp recombinase under control of the endogenous Kirrel3 gene. A Flp sequence was inserted after the start codon of Kirrel3 (Figure 1A) along with a viral 2A sequence to allow for normal expression of the Kirrel3 protein (Figure 1B). In the hippocampus, Kirrel3 is expressed by DG neurons and some GABA neurons (Martin, Muralidhar et al. 2015). To test the specificity of our new mouse line, we crossed the Kirrel3-Flp line to a Gad2-Cre line expressing Cre in all GABA neurons (Taniguchi, He et al. 2011) and to a reporter line known as RC:FLTG (Plummer, Evsyukova et al. 2015). This reporter labels cells co-expressing both Flp and Cre with GFP (K3-GABA neurons) and cells expressing Flp alone with tdTomato (DG neurons) as predicted (Figure 1C). Next, we verified that GFP-positive cells are in fact K3-GABA neurons. Because there is no suitable antibody to label Kirrel3 protein in tissue, we used in situ hybridization with mRNA probes for Kirrel3, GAD1, and GFP (Figure 1D). We imaged cells in area CA3 and found that the transgenic line functions as expected. 86% ± 17 (mean ± SEM) of GFP cells express Kirrel3 (Figure 1E), indicating that GFP expression marks Kirrel3 cells. Moreover, 94% ± 14 (mean ± SEM) of Kirrel3 cells in area CA3 co-express GAD1 and, therefore, are likely GABA neurons (Figure 1E). This experiment also indicates that ∼34% ± 14 (mean ± SEM) of all GABA neurons express Kirrel3 mRNA (Figure 1E). This is slightly more than previously reported (∼19%) using immunohistochemistry on Kirrel3 knockout mice (Martin, Muralidhar et al. 2015), which express GFP instead of Kirrel3. The different percentages are likely due to different sensitivities of each method and age of the animals.

**Figure 1.**
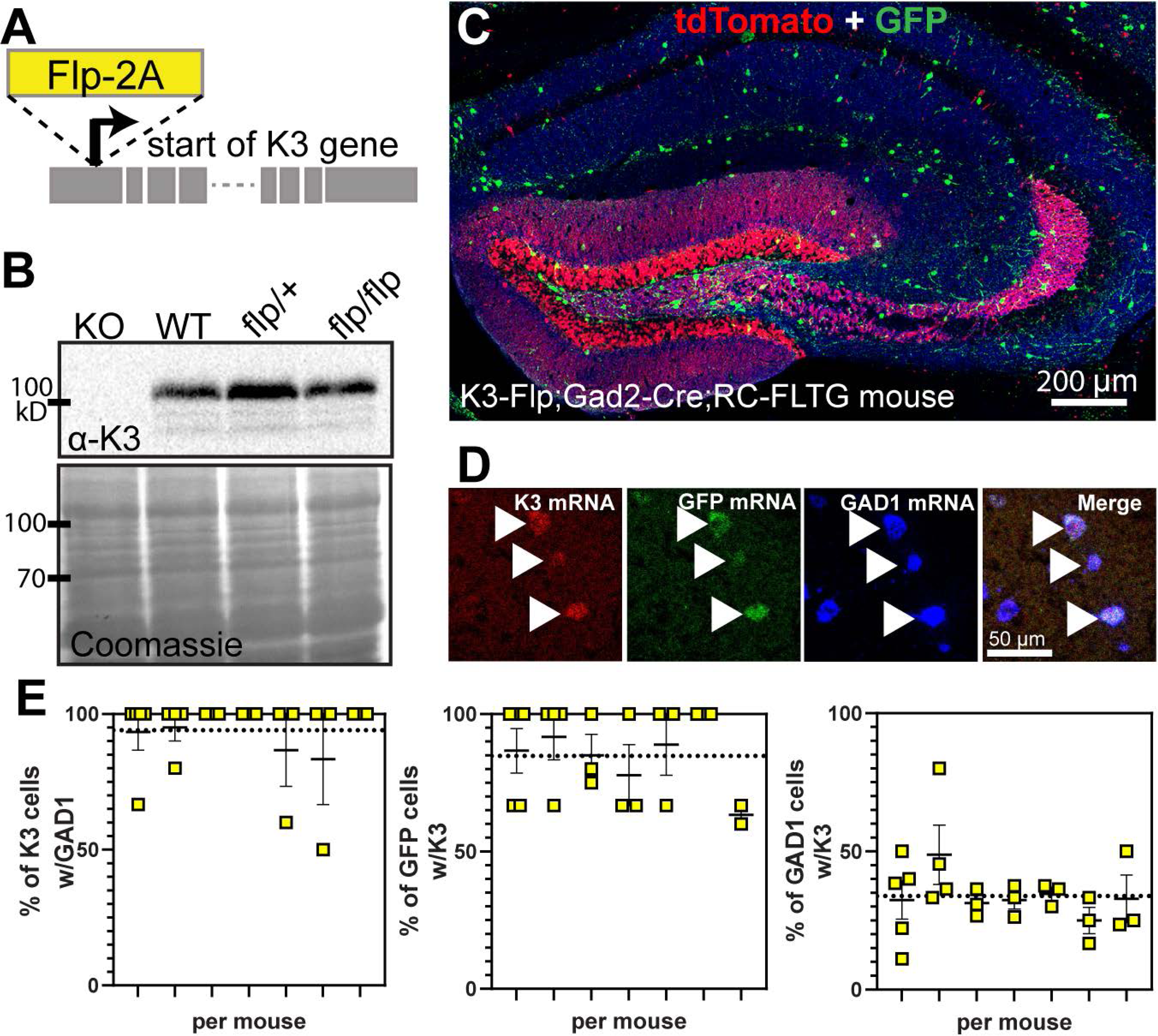
Generation and validation of a Kirrel3-Flp driver mouse line. A) Schematic of the Flp-2A insertion site in the mouse Kirrel3 gene. B) Western blot of brain lysates from 2 months old mice show that the Flp transgenic line expresses normal levels of Kirrel3 protein. Lysates from Kirrel3 knockout (KO) and wildtype (WT) are used as controls. Coomassie stained membrane indicates equal protein loading. C) A hippocampal section from an adult Kirrel3-Flp;Gad2-Cre;RC-FLTG triple transgenic mouse (heterozygous for all transgenes indicated) stained with antibodies against TdTomato (red) and GFP (green). D) A magnified image of a hippocampal section labeled by fluorescent in situ hybridization (FISH). Arrows indicate K3-GABA neurons co-expressing Kirrel3, GFP, and GAD1 mRNA. E) Quantification of FISH as indicated. Each column represents a mouse and each square represents one hippocampal section from that mouse. Average percentages from all data points are shown with a dotted line. n = 7 adult mice (3 male and 4 female) Error bars represent SEM.

To study K3-GABA neuron function in vivo, we used a newly developed intersectional AAV that expresses the activating DREADD hM3D in a Cre and Flp-dependent manner. This strategy, called “ConVERGD” (Hughes, Pittman et al. 2024), contains a LoxP-flanked ribozyme, which must be removed by Cre to prevent mRNA degradation, and an inverted, Frt-flanked activating DREADD hM3D-mCherry payload driven by the human synapsin promoter (Figure 2A) (Hughes, Pittman et al. 2024). We injected the hM3D-mCherry ConVERGD virus into dorsal hippocampal area CA3 of K3-Flp;Gad2-Cre double heterozygous mice to specifically express hM3D-mCherry in K3-GABA neurons. We observed cell labeling in a pattern consistent with the location of K3-GABA neurons with no off-target expression in Kirrel3-Flp or wildtype mice (Figure 2B). We also used a dual FISH/immunostaining protocol to confirm that the hM3D-mCherry protein is expressed in cells that co-express Kirrel3 and Gad1 mRNA (K3-GABA neurons) (Figure 2C). Finally, we tested if hM3D-mCherry expression increases the activity of K3-GABA neurons in the brain in a ligand-dependent manner in vivo. For this experiment, we injected hM3D-mCherry ConVERGD virus into dorsal CA3 of K3-Flp;Gad2-Cre mice, allowed the virus to express for 3 weeks, and immunostained for cFos 40 minutes after a foot shock administration (Figure 2D). We found that DCZ administration 10 minutes prior to foot shock significantly increased the percentage of K3-GABA neurons expressing cFos compared to saline injection (Figure 2E, F). Our results demonstrate that we can selectively label and activate K3-GABA neurons in vivo to study their function in the mouse brain.

**Figure 2:**
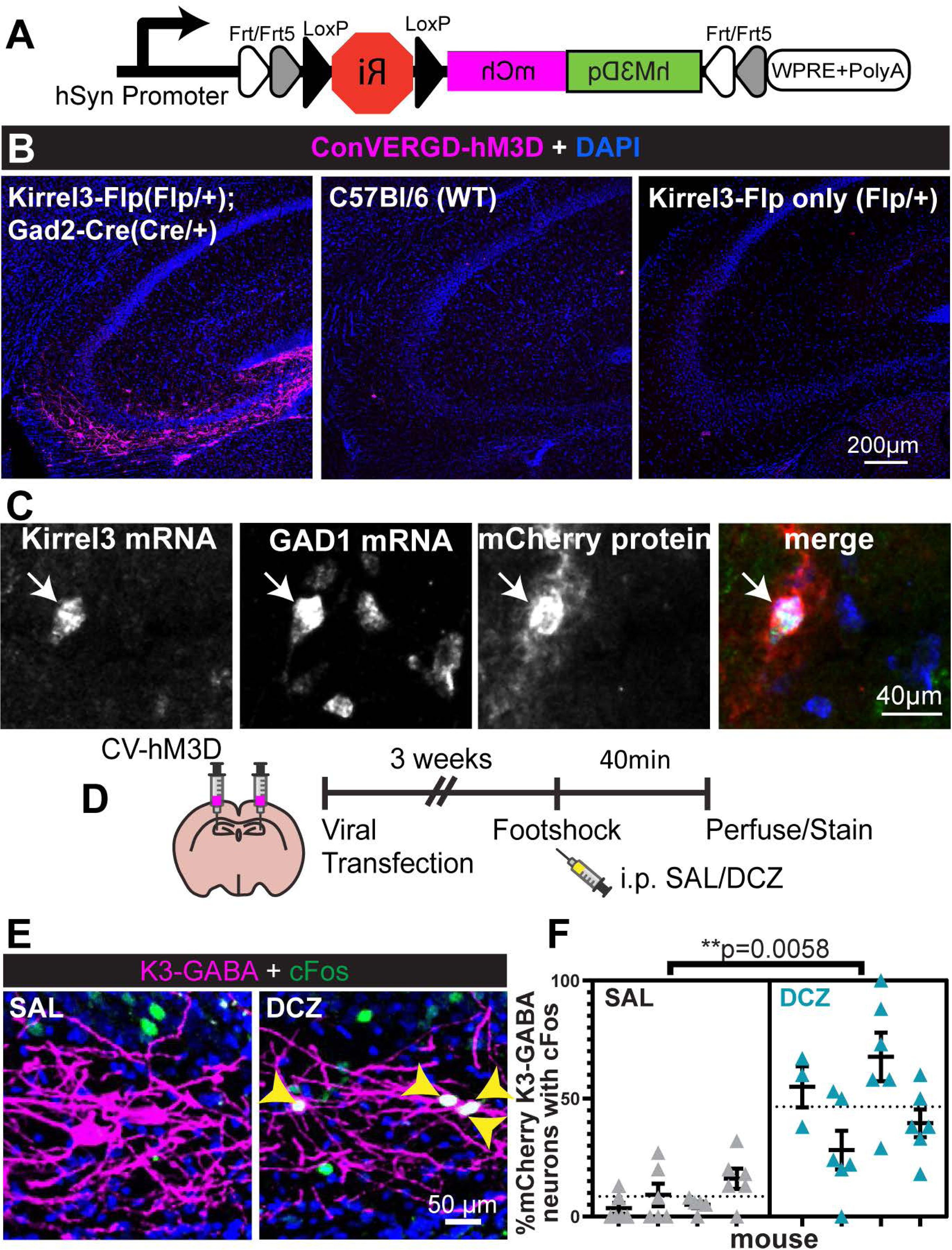
Intersectional genetics enables specific targeting and activation of K3-GABA neurons. A) Schematic of the ConVERGD (CV)-hM3D-mCherry AAV construct. B) Representative hippocampal images from an adult mice (genotypes indicated) infected with CV-hM3D-mCherry targeted to area CA3. mCherry is only present when both Flp and Cre are expressed (left). C) Magnified image from a hippocampal section processed for dual FISH/immunohistochemistry from a Kirrel3-Flp;Gad2-Cre mouse infected with CV-hM3D-mCherry. D) Schematic of the experimental design for E and F. E) Representative images showing cFos (green) in K3-GABA neurons expressing hM3D-mCherry (magenta) specifically after DCZ injection (right). Yellow arrowheads indicate K3-GABA neurons that express cFos (merge is white). F) Quantification of the % of DREADD-expressing K3-GABA neurons that express cFos for saline and DCZ treated mice after foot shock. n = 4 males each. Each column represents one mouse, up to 3 sections and 6 hippocampi counted per mouse. Average from all data points is shown with a dotted line. Error bars represent SEM, nested t-test.

### Activating K3-GABA neurons impairs memory discrimination

Given that hippocampal circuits are critically important for learning and memory, we sought to determine if and how activating K3-GABA neurons alters learning and memory in mice. To do this, we adapted a contextual fear conditioning paradigm that measures context learning, recall, and discrimination across multiple contexts (Figure 3A, Extended Data Figure 3-1) (Nakashiba, Cushman et al. 2012, Sun, Bernstein et al. 2020). We conditioned the mice in context A where they received 4 unsignaled foot shocks (1.0 mA, 2 sec duration, 60 sec inter-stimulus-interval) (Figure 3A). One day later, we tested their fear memory by measuring freezing behavior when they were placed back in context A without foot shock (Figure 3A). On the same day, we also measured freezing behavior in a similar but slightly different context B to test their ability to discriminate between two closely related contexts (test order counterbalanced) (Figure 3A). We repeated the procedure the next day but this time we tested if mice discriminate between the conditioned context A and a novel, very distinct context C (order counterbalanced) (Figure 3A). We quantified their ability to discriminate between the two contexts each day by calculating a discrimination ratio (A/A+B) or (A/A+C) in which a value of 1.0 indicates the mouse only exhibited freezing behavior in context A, a value of 0.5 indicates the mouse exhibited equal freezing behavior in both contexts, and a value of 0 would indicate the mouse only exhibited freezing behavior in context B or C (Poulos, Mehta et al. 2016).

**Figure 3.**
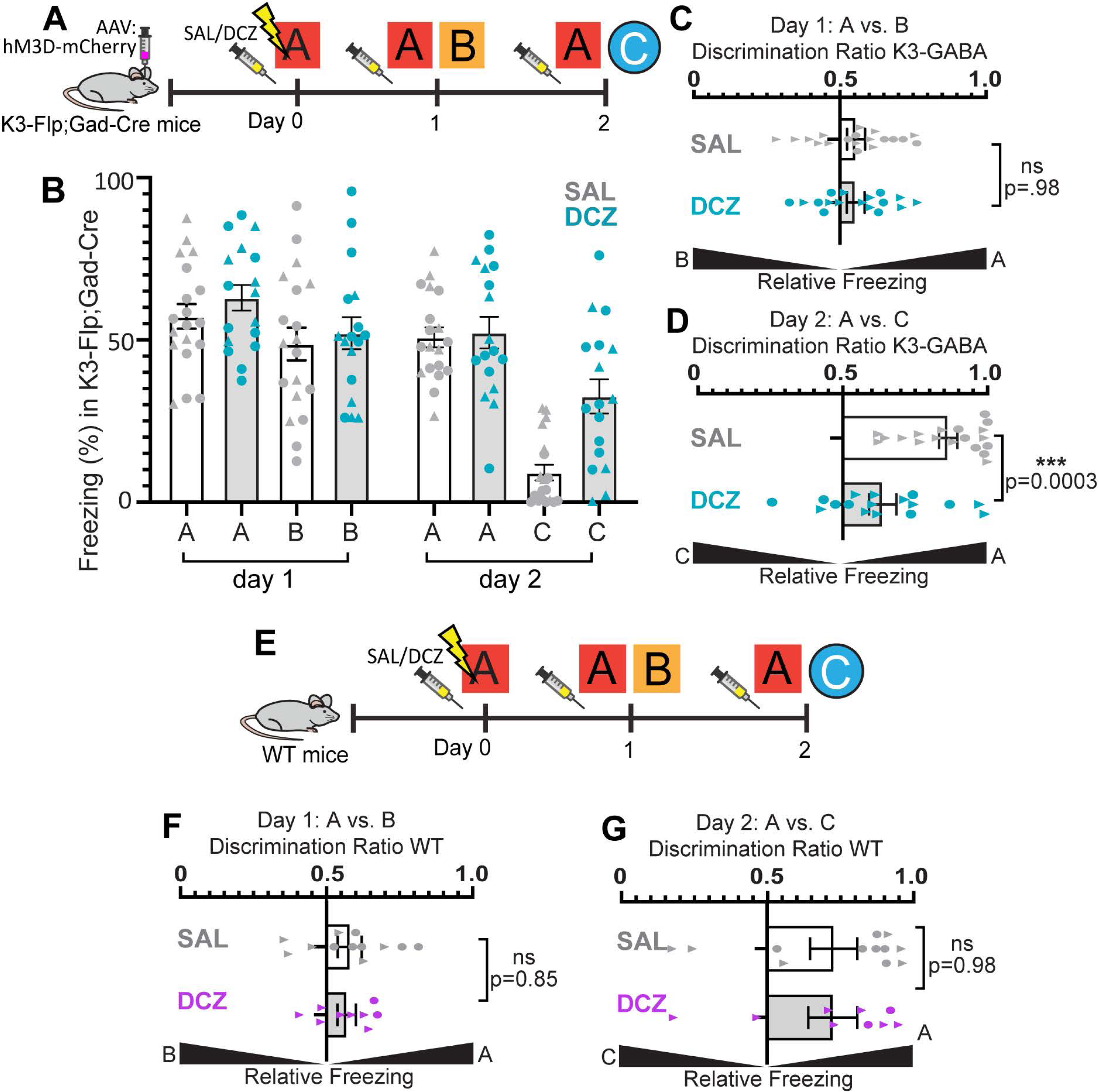
K3-GABA activation impairs memory discrimination. A) Schematic of behavioral paradigm and experimental design for K3-Flp;Gad2-Cre mice. B) Time spent freezing (%) when placed in indicated contexts after saline (gray) or DCZ (blue) injection. All mice were adult K3-Flp;Gad2-Cre heterozygotes injected with CV-hM3D-mCherry. n = 19 saline, 17 DCZ mice. C) Context discrimination ratios for K3-GABA mice on day 1 (A vs B). mean +/-SEM, unpaired t-test. D) Context discrimination ratios for K3-GABA mice on day 2 (A vs C). mean +/-SEM, unpaired t-test. E) Experimental design for wildtype (WT) no DREADD control mice. F, G) Discrimination ratios from WT mice for contexts A vs B (F) or contexts A vs C (G). n = 12 saline; 9 DCZ. Mean +/-SEM, unpaired t-test. In all graphs, males are represented with a triangle, females a circle.

To determine if activating K3-GABA neurons affects the learning, memory, or discrimination of contextual fear memories, K3-Flp;Gad2-cre heterozygous mice were bilaterally infected with the ConVERGD hM3D-mCherry AAV in dorsal hippocampal area CA3 (Figure 3A). Because we do not know whether K3-GABA neurons participate in any aspect of memory acquisition or recall, we initially opted to give saline or DCZ injections 10 minutes prior to every context placement (Figure 3A). Regardless of treatment, all mice were able to mount activity bursts in response to foot shock on training day 0, showing no difference in context fear acquisition (Extended Data Figure 3-1) and no differences in baseline freezing or activity before the shocks were given (Extended Data Figure 3-1), indicating that K3-GABA neuron activation does not cause overt changes to motor control.

Consistent with prior work (Lovett-Barron, Kaifosh et al. 2014, Besnard and Sahay 2016, Bernstein, Does et al. 2021), we found that control saline-injected mice show poor discrimination between very similar contexts, exhibited by a similarly high rate of freezing behavior in A and B (Figure 3B and C) and an average discrimination ratio near 0.5. Moreover, as expected, control mice, on average, show robust discrimination between the very distinct contexts A and C (Figure 3B and D). Activating K3-GABA neurons did not significantly change the fear response to the conditioning context A or the similar context B (Figure 3B). However, activating K3-GABA neurons robustly increased the fear response in the neutral context C on day 2 (Figure 3B). This resulted in a significant decrease in the average A/C discrimination ratio compared to saline controls (Figure 3D). We confirmed proper hM3D-mCherry expression and targeting to the CA3/hilus region in all mice used for experiments (Extended Data Figure 3-1). We verified that the change in behavior was not due to DCZ itself by testing mice that do not express the DREADD receptor in our behavior paradigm (Figure 3E). DCZ-injected wildtype mice that do not express the DREADD receptor showed similar fear responses and discrimination ratios to saline controls (Figure 3F,G and Extended Data Figure 3-1). Thus, our data indicates that activating K3-GABA neurons located in the CA3/hilus region erodes memory precision by causing mice to overgeneralize fear to a distinct, neutral context.

### Activating PV neurons does not impair memory discrimination

To test if K3-GABA-mediated deficits in memory discrimination are specific to K3-GABA neurons or could be attributed to the activation of any similarly sized subset of GABA neurons, we sought to identify an alternative population of GABA neurons. We previously noted that K3-GABA neurons are heterogeneous and the majority of them do not fall into any one commonly studied, molecularly-defined subgroup of GABA neurons (Martin, Muralidhar et al. 2015). We revisited this idea by immunostaining K3-GABA neurons labeled using our newly developed Kirrel3-Flp reporter line with common GABA subtype markers (Extended Data Figure 4-1) and by searching single cell sequencing databases (Yao, van Velthoven et al. 2021). As before, we found that K3-GABA neurons are transcriptionally heterogenous, but that they consistently have little overlap with PV neurons (Extended Data Figure 4-1) (Martin, Muralidhar et al. 2015), which are a similarly sized population of hippocampal GABA neurons (Fukuda, Heizmann et al. 1997, Gradwell, Boyle et al. 2022). Thus, we tested if activation of PV neurons impairs memory discrimination using the same behavioral paradigm. To activate PV neurons, we injected PV-Cre mice with a Cre-dependent hM3D-mCherry (Krashes, Koda et al. 2011). As expected, PV immunostaining revealed that Cre-driven hM3D-mCherry expression was restricted to neurons that express PV (Extended Data Figure S2C). As we did for K3-GABA neurons, we tested if DCZ treatment significantly increases cFos expression in PV neurons expressing hM3D after foot shock compared to saline control treatment. We found that PV neurons expressing hM3D are significantly activated by injection of DCZ (Figure 4A, B). Next, we tested if DREADD activation of PV neurons alters memory discrimination using the context fear conditioning paradigm (Figure 4C). Unlike our results for K3-GABA neurons, we found that activating PV neurons with DCZ does not alter memory discrimination when compared to saline controls (Figure 3D-F). We confirmed that PV-Cre mice had correct viral expression and that fear acquisition and baseline activity were normal (Extended Data Figure 4-1). Our results indicate that K3-GABA neurons in area CA3 have a specific function in memory discrimination that is not shared with PV neurons.

**Figure 4.**
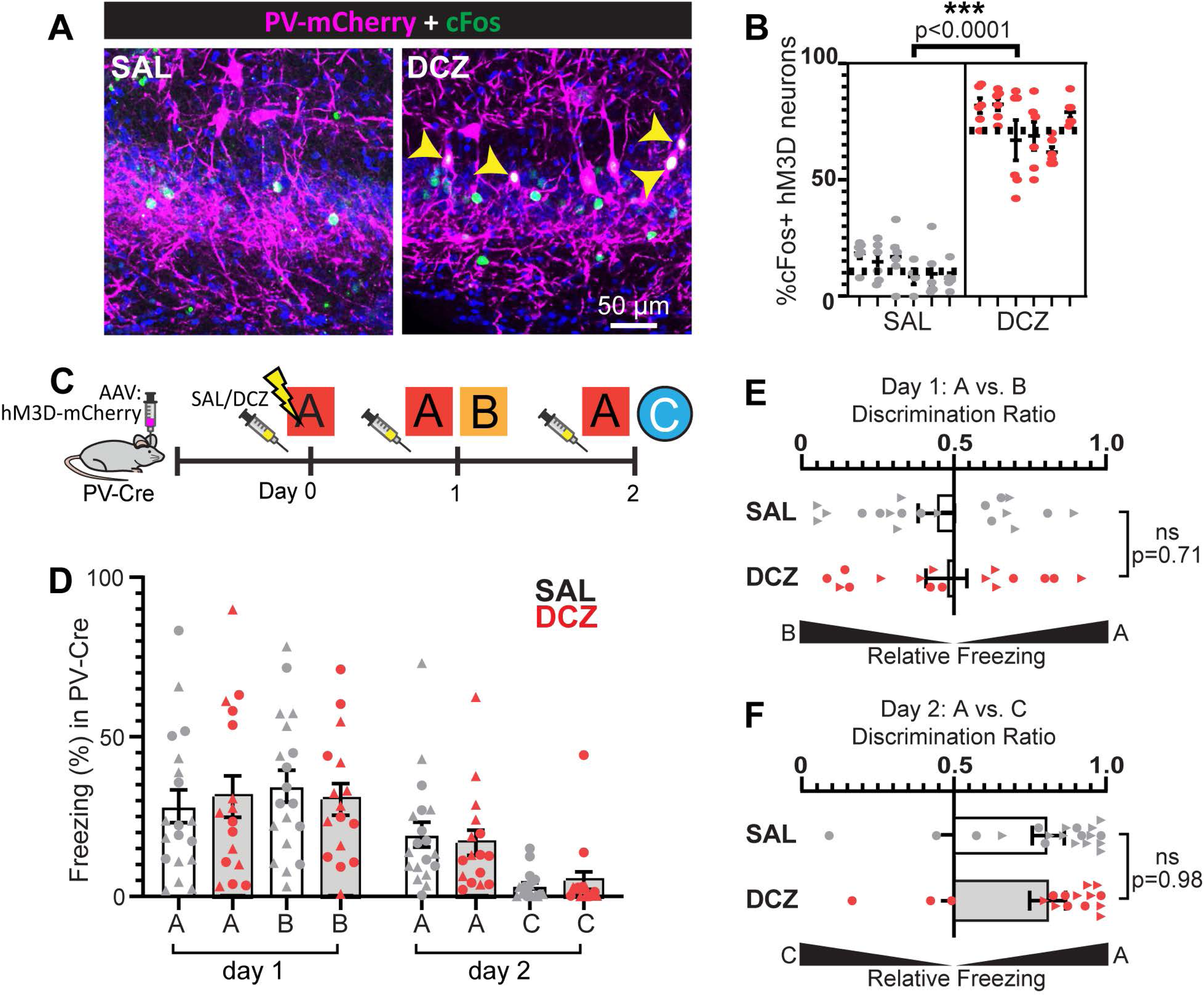
Activating parvalbumin-expressing neurons does not impair memory discrimination. A) Example sections from adult PV-Cre mice expressing the Cre-dependent DIO-hM3D-mCherry AAV and stained for cFos (green). B) % of hM3D-expressing PV neurons that are cFos positive after saline (gray) and DCZ injection (red) following foot shock. n= 6 mice each condition, global average indicated by dotted line, nested t-test. C) Experimental design for PV-Cre mice. D) Time spent freezing (%) when placed in indicated contexts after saline (gray) or DCZ (red) injection. All mice were adult PV-Cre heterozygous mice injected with Cre-dependent hM3D-mCherry. n= 19 saline, 16 DCZ. E,F) Data from D plotted as discrimination ratios for contexts A vs B (E) or contexts A vs C (F). Error bars represent SEM, t-tests. Each data point represents a mouse with males a triangle, females a circle.

### Activating K3-GABA neurons during recall impairs memory discrimination

Given that global activation of K3-GABA neurons across all phases of fear conditioning selectively impaired memory discrimination between two distinct contexts (Figure 3), we next tested if this effect was primarily caused by K3-GABA activation during context conditioning or during context recall. To test this, K3-Flp;Gad2-cre heterozygous mice expressing hM3D-mCherry in area CA3 K3-GABA neurons were subject to our three-context fear conditioning task. This time, mice received saline or DCZ injections either prior to foot shock conditioning in context A (Figure 5A) or prior to the presentation of contexts A and C on day 2 (Figure 5E). As before, all mice showed normal baseline activity and fear acquisition to the foot shock (Extended Data Figure 5-1). Interestingly, activation of K3-GABA neurons during conditioning had no effect on fear responses in any of the recall contexts on days 1 or 2 and these mice showed normal discrimination between the fear context A and the neutral context C (Figure 5B-D). However, activation of K3-GABA neurons only during day 2 recall reproduced the memory discrimination phenotype that we observed when K3-GABA neurons were activated at all time points. Mice in which K3-GABA neurons were selectively activated only on day 2 showed significantly worse memory discrimination between the fear context A and the neutral context C (Figure 5F-H). Together, these results suggest that K3-GABA neuron activity levels do not modulate contextual encoding but that they are critical for memory discrimination during recall, suggesting K3-GABA neurons gate the ability of animals to retrieve memories with precision.

**Figure 5.**
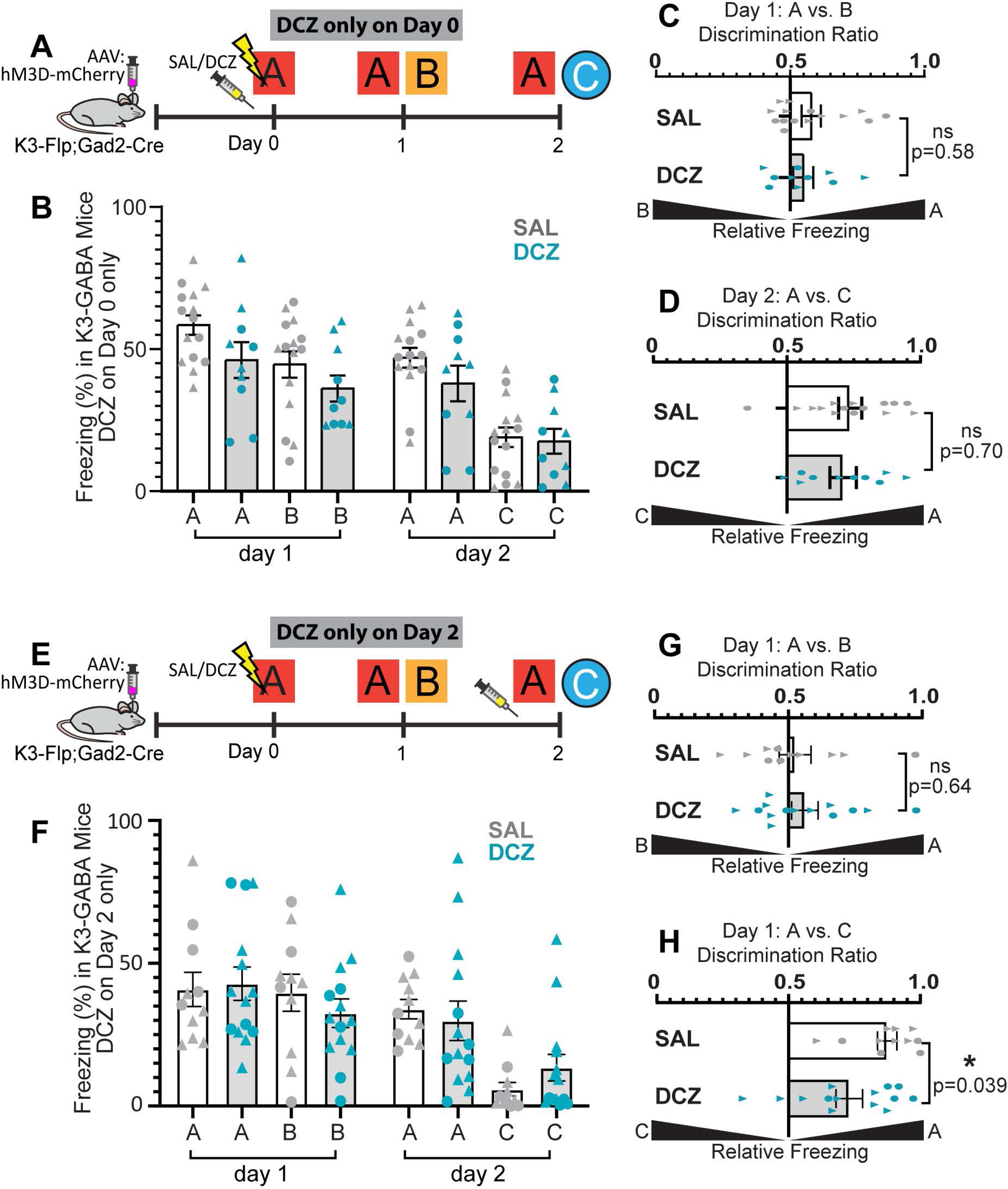
K3-GABA activation during recall impairs memory discrimination. A) Experimental design showing saline or DCZ injection only prior to foot shock conditioning in context A in K3-Flp;Gad2-Cre heterozygotes mice. B) Mouse behavior plotted as percent time spent freezing when placed in indicated contexts. Here, DCZ refers to the mice that received DCZ at conditioning. They were not treated with DCZ prior to each context. n= 15 saline, 10 DCZ mice. C,D) Data from B plotted as discrimination ratios for contexts A vs B (C) or contexts A vs C (D). Error bars indicate SEM, t-tests. E) Experimental design showing saline or DCZ injection only prior to contexts A and C on day 2 in K3-Flp;Gad2-Cre heterozygotes mice. F) Mouse behavior plotted as percent time spent freezing for when placed in indicated contexts. Here, DCZ refers to the mice that received DCZ on day 2 only. They were not treated with DCZ on other days. n= 11 saline, 14 DCZ mice. G,H) Data from F plotted as discrimination ratios for contexts A vs B (G) or contexts A vs C (H). Error bars indicate SEM, t-tests. In all graphs, each data point represents a mouse with males a triangle, females a circle.

### K3-GABA neuron activity strongly inhibits CA3 neuron activity

To understand how K3-GABA neurons uniquely modulate hippocampal circuits compared to PV neurons, we performed electrophysiological recordings from CA3 in awake, behaving mice (Figure 6A). We virally expressed hM3D-mCherry in either K3-GABA or PV neurons of adult mice, and five days later, mice began training to run on a cylindrical treadmill through a 3-meter linear virtual environment. After training, we implanted a Neuropixels probe targeting CA3.

**Figure 6.**
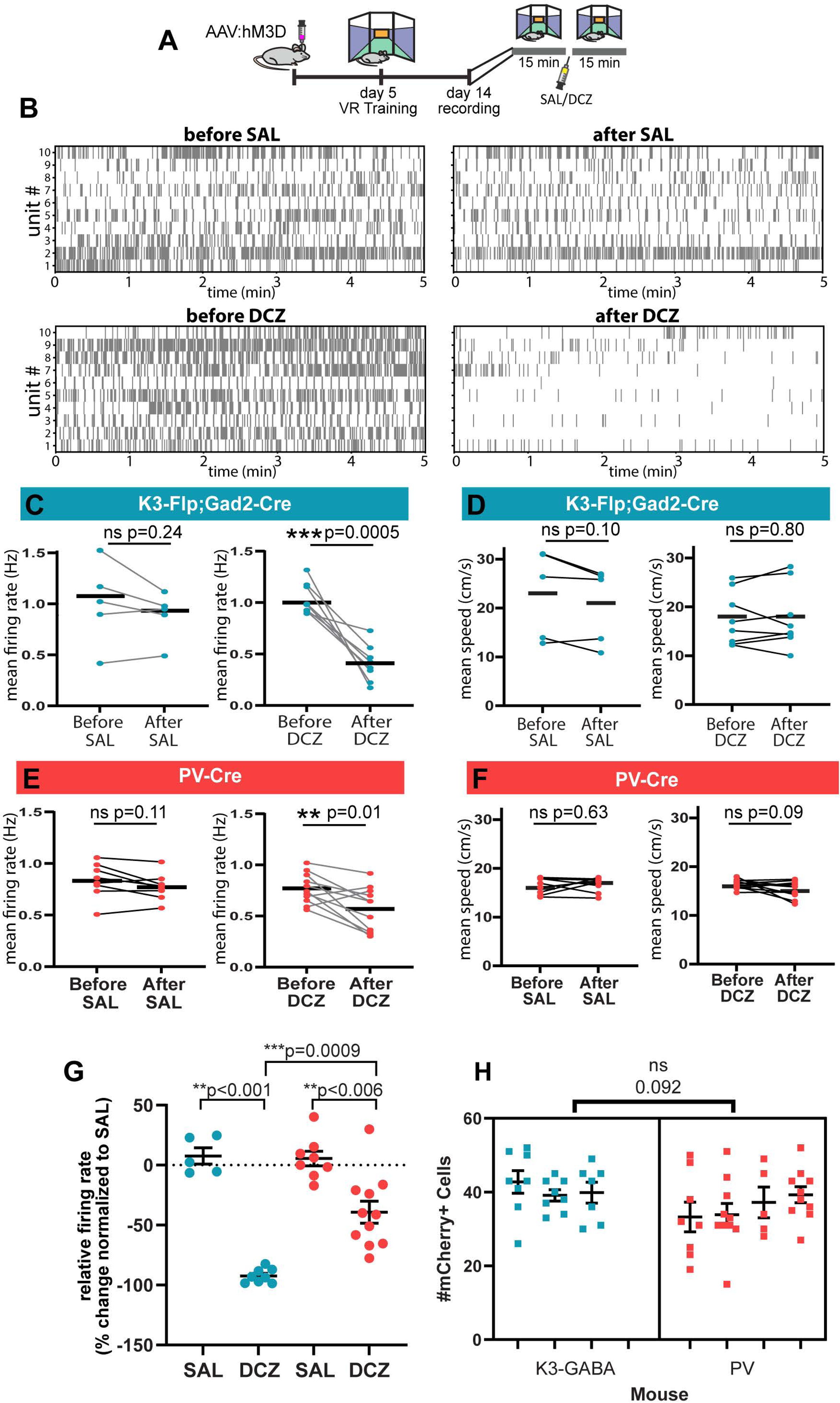
K3-GABA neuron activity strongly inhibits CA3 neuron spike rate. A) Experimental design for in vivo recordings. B) Raster plots of representative single unit CA3 recordings from mice before and after saline or DCZ injection. 10 units each graph. C) Mean firing rate of CA3 single units for each recording session for hM3D-mCherry infected K3-Flp;Gad2-Cre mice treated with saline or DCZ. n=5 saline; 8 DCZ, paired t-test. D) Mean running speed for hM3D-mCherry infected K3-Flp;Gad2-Cre mice treated with saline or DCZ. n=5 saline; 8 DCZ, paired t-test. E) Mean firing rate of CA3 single units for each recording session for hM3D-mCherry infected PV-Cre mice treated with saline or DCZ. n=8 saline; 11 DCZ, paired t-test. F) Mean running speed for hM3D-mCherry infected PV-Cre mice treated with saline or DCZ. n=8 saline; 11 DCZ, paired t-test. G) Firing rates were normalized as a percent change from saline for each genotype. Graph summarizing and comparing data from C and E. ANOVA with post-tests. H) Average number of DREADD expressing neurons per section from each mouse used for in vivo recording experiments in B-G. nested t-test.

Using the VR setup and treadmill velocity measurements, we were able to rule out differences in running speed as a confounding factor in firing rate comparisons across control and experimental conditions during DREADD-mediated manipulations. After probe insertion and a mock intraperitoneal injection, a baseline 15-minute recording was performed on the virtual track, followed by an injection of either saline or DCZ and another 15-minute recording (Figure 6A). We obtained between 50-100 or more single unit recordings in CA3 for each session of each mouse. Figure 6B shows spiking activity from 10 representative single unit CA3 recordings from two K3-GABA mice, one given saline and one given DCZ. It was apparent that CA3 spiking activity is strongly and significantly reduced after DCZ injection in mice in which K3-GABA neurons are activated (Figure 6B and C). In contrast, running speed, which can affect firing rates, did not change before or after DCZ or saline injection (Figure 6D). Activating PV neurons also decreased CA3 activity without affecting running speed (Figure 6E and F). This is not unexpected because PV neurons are inhibitory. However, activation of K3-GABA neurons decreased CA3 activity to a significantly greater extent than activation of PV neurons when directly compared (Figure 6G). We verified that the observed differences were not simply due to differences in number of DREADD-expressing cells in each genotype as the number of infected cells was very similar among all mice tested (Figure 6H). This indicates that K3-GABA neuron activity specifically exerts powerful control over CA3 spiking activity.

### K3-GABA neurons act locally within the CA3

We next considered that K3-GABA neurons located in the hilus and CA3 could inhibit CA3 pyramidal activity via direct innervation or indirectly by projecting back to the DG, which is an important source of excitation to CA3 (Figure 7A). To investigate this, we used cFos immunostaining to examine neural activity in different populations of principal neurons throughout the hippocampus after we chemogenetically activated K3-GABA neurons. For this experiment, mice expressing hM3D-mCherry in K3-GABA neurons located in the hilus/CA3 area were injected with saline or DCZ and then placed in a novel enriched environment to broadly stimulate neuronal activity (Figure 7B). We then assessed the density of cFos expression in K3-GABA neurons and each hippocampal subregion using immunohistochemistry. As expected, DCZ treatment resulted in a significant increase in the number of cFos-positive K3-GABA neurons compared to saline (Figure 7C). This is consistent with our prior experiment using foot shock (Figure 2E,F) and indicates that the DREADD works as expected to activate K3-GABA neurons in vivo. We also found that activating K3-GABA neurons with DCZ results in a significant reduction of cFos-positive neurons in CA3 compared to saline-injected mice (Figure 7D). This is consistent with the in vivo electrophysiology and indicates that activation of K3-GABA neurons inhibits CA3 pyramidal neuron activity. However, we found no significant changes in the activity of neurons in the DG, CA1, or hilus regions after K3-GABA neuron activation (Figure 7E-G). These results strongly suggest that K3-GABA neurons directly inhibit CA3 neurons and do not act indirectly through the DG. Interestingly, activating PV neurons (Figure 7H) does not change the density of active CA3 neurons as measured by cFos (Figure 7I). Although we just reported that activating both K3-GABA and PV neurons inhibits CA3 spike rates, K3-GABAs suppress CA3 spiking to a significantly greater extent (Figure 6G). Our cFos analysis suggests that this difference in spike rate may be functionally relevant because, at the transcriptional level of immediate early gene expression, Kirrel3-GABA neuron activation significantly suppresses cFos activity in CA3 neurons, but PV-GABA neuron activation does not.

**Figure 7.**
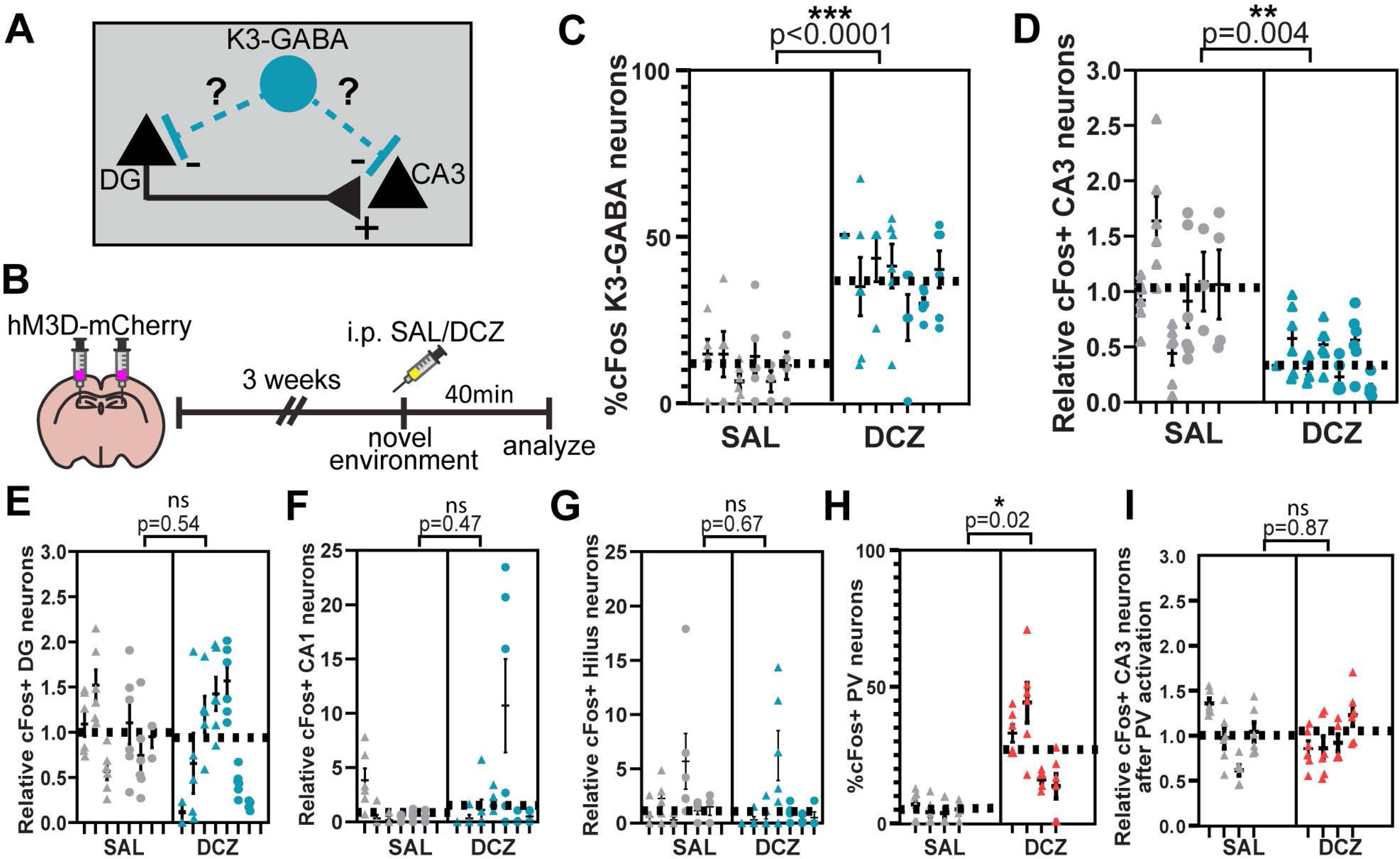
Activating K3-GABA neurons specifically decreases cFos expression in area CA3. A) Schematic of DG-to-CA3 connectivity and the potential connectivity of K3-GABA neurons to mediate either direct inhibition to CA3 or indirect inhibition via DG. B) Experimental design for C-I. C) Percent of DREADD-expressing K3-GABA neurons that are cFos positive. n=6 saline, 7 DCZ mice. nested t-test. D-G) Relative density (normalized to saline controls) of cFos-positive cells in CA3, DG, CA1, and hilus in hM3D-mCherry infected K3-Flp;Gad2-Cre mice treated with saline or DCZ. n=6 saline, 7 DCZ mice. nested t-test. H) Percent of DREADD-expressing PV neurons that are cFos positive. n=4 saline, 4 DCZ mice. nested t-test. I) Relative density (normalized to saline controls) of cFos-positive cells in CA3 in hM3D-mCherry infected PV-Cre mice treated with saline or DCZ. n=4 saline, 4 DCZ mice. nested t-test. For all graphs, each point represents a hippocampus section, each column represents a mouse, males a triangle, females a circle, and the average for all data points in a condition is shown as a dotted line. In all graphs, blue data points indicate analyses done in mice with K3-GABA neurons activated and red data points indicate mice with PV neurons activated. Error bars show SEM.

### K3-GABA neurons receive direct input from DG mossy fiber boutons and synapse onto CA3 dendrites

Finally, to gain more information about how K3-GABA neurons inhibit CA3, we investigated the synaptic connectivity of K3-GABA neurons. Prior work indicated that Kirrel3 knockout mice selectively lose DG-to-GABA mossy fiber filopodia synapses in the mouse brain (Martin, Muralidhar et al. 2015, Martin, Woodruff et al. 2017). These filopodia synapses are one part of the large mossy fiber synaptic complex made in DG axons. Here, the main mossy fiber bouton is a huge glutamatergic “detonator” that releases neurotransmitters at many active zones with a single CA3 neuron (Rollenhagen and Lübke 2010, Marosi, Arszovszki et al. 2023). During development, DG axons sprout filopodial projections directly off the main bouton that synapse with nearby GABA neurons and these are thought to provide feed-forward inhibition to CA3 (Torborg, Nakashiba et al. 2010, Guo, Soden et al. 2018). Because Kirrel3 is a homophilic cell adhesion molecule that promotes synapse formation in vitro (Taylor, Martin et al. 2020), it strongly suggested that mossy fiber filopodia specifically synapse onto Kirrel3-positive GABA neurons, but it could not be easily tested until we made the Kirrel3-Flp line described here.

Mossy fiber filopodia are easily visualized with light microscopy but they are transient structures that are largely absent in adults (Wilke, Antonios et al. 2013). Therefore, to test if K3-GABA neurons receive mossy fiber input, we used electron microscopy (EM). We labeled K3-GABA neurons with GFP in K3-Flp;Gad2-Cre heterozygous mice with an intersectional GFP AAV that can only be expressed in neurons co-expressing Flp and Cre (Fenno, Mattis et al. 2014). We then correlated light and EM images to identify a K3-GABA neuron cell body before using serial blockface scanning EM to generate a high-resolution EM dataset in the identified area (Figure 8A-C). We reconstructed several dendritic branches, including one that projected directly into the mossy fiber layer (Figure 8D-F). This dendritic branch measures 73 μm and extends 34 μm in the stratum lucidum layer that houses mossy fiber synapses (Figure 8D-F). Here it is directly innervated by at least 14 different mossy fiber boutons with some boutons making more than one synapse onto the K3-GABA dendrite (Figure 8E and G) and two boutons make filopodia synapses onto the K3-GABA dendrite. The K3-GABA dendrite is also innervated by over 25 en passant synapses (Figure 8F and H). Most (16/25) of the en passant synapses are from axons traveling in the same plane as mossy fibers and are most likely also from DG axons. Thus, our results strongly suggest that DG mossy fibers directly innervate K3-GABA neurons using both large mossy fiber boutons and more typical en passant synapses.

**Figure 8.**
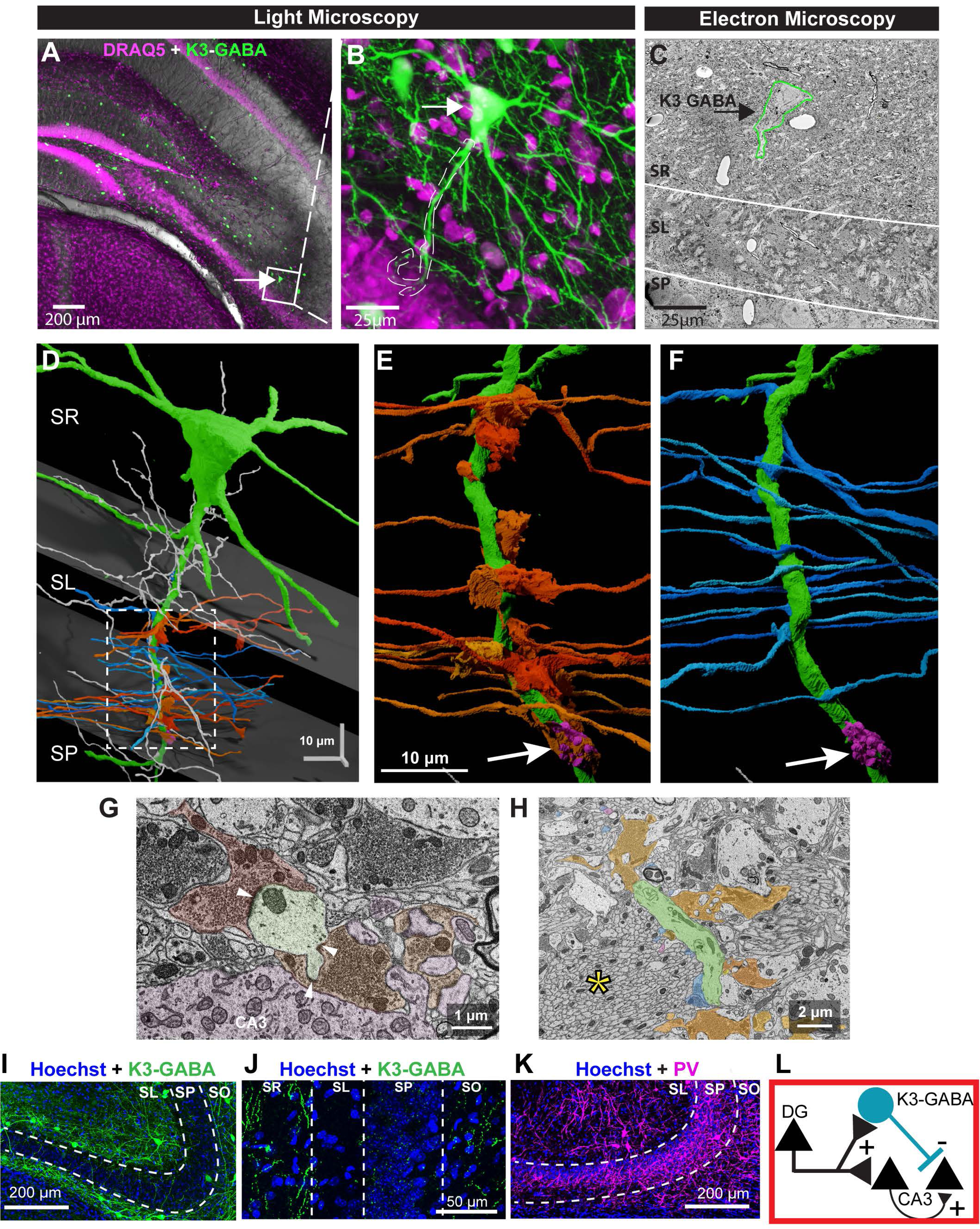
K3-GABA neurons receive direct input from DG mossy fiber boutons. A,B) Hippocampal section from an 84 days old Kirrel3-Flp;Gad2-Cre mouse injected with an AAV that mediates expression of a Flp and Cre-dependent GFP. Kirrel3-GABA neurons are green, the DRAQ5 nuclear stain is magenta. The arrow points to a Kirrel3-GABA neuron shown magnified in B. C) The same Kirrel3-GABA neuron identified in B is outlined in green in an EM view. D) A 3D reconstruction of the identified K3-GABA dendrite (green). Gray planes note the top and bottom of the SL layer. DG axons with mossy fiber boutons that synapse onto the Kirrel3-GABA dendrite are orange, DG axons with en passant synapses are blue and other axons are gray. E,F) A magnified view of the Kirrel3-GABA dendrite and innervating axons. Colors are the same as in D. The arrow points to a CA3 thorny excrescence spine (pink) G) EM image of the Kirrel3-GABA dendrite (green) receiving multiple inputs from two mossy fiber boutons (orange). A nearby CA3 neuron and its associated multi-headed thorny excrescence spine are shown (pink). Arrowheads indicate synaptic densities from mossy fiber boutons to the Kirrel3-GABA dendrite. H) A less magnified EM image showing axons connected via large mossy boutons (orange) and en passant synapses (blue). Large bundles of DG axons can be seen (asterisk). I) Hippocampal section from an adult Kirrel3-Flp;Gad2-Cre mouse infected with a Flp and Cre-dependent GFP AAV. K3-GABA neurons express GFP (green), Hoechst nuclear stain (blue). J) Higher magnification image of a Kirrel3-Flp;Gad2-Cre mouse sparsely infected with a Flp and Cre-dependent GFP AAV. Note Kirrel3-GABA axon labeling primarily in the SR and SO layers. Hoechst nuclear stain (blue). K) Hippocampal section from a PV-Cre mouse infected with a Cre-dependent AAV driving expression of mCherry. PV neurons express mCherry (magenta) and Hoechst nuclear stain (blue). SO; stratum oriens, SP; stratum pyramidale, SL; stratum lucidum, SR; stratum radiatum

Unfortunately, the axon of the identified K3-GABA neuron quickly left the EM field of view, and we could not track the location of its output synapses. Therefore, we instead bulk labeled K3-GABA neurons, including their axons, with a Flp and Cre-dependent GFP reporter (Figure 8I). Many K3-GABA dendrites and cell bodies reside in the SL and SO layers (Figure 8I), and a magnified image from a sparsely transfected area lacking transfected cell bodies indicates that K3-GABA axons are primarily located in SR and SO layers (Figure 8J), which are the sites of recurrent CA3-CA3 collateral synapses. This strongly suggests K3-GABA neurons target CA3 pyramidal cell dendrites and are positioned to inhibit recurrent CA3 activity. For comparison, we also labeled PV neurons in area CA3 by injecting a Cre-dependent mCherry reporter into PV-Cre mice. We noted that PV neurons have the opposite pattern with extensive arborizations in the CA3 pyramidal cell body layer (Figure 8K), suggesting they target CA3 cell bodies as expected. Taken together, our data indicates that at least some K3-GABA neurons receive input from DG and suggests that most K3-GABA neurons likely mediate feed-forward inhibition onto CA3 neuron dendrites.

## DISCUSSION

Here, we identified a population of GABA neurons based on expression of the synaptic cell adhesion molecule Kirrel3. We demonstrate that increasing the activity of K3-GABA neurons has two main effects. On the cellular level, K3-GABA activity robustly inhibits CA3 neuron activity and at the behavioral level, K3-GABA activity decreases memory discrimination. A reduction in memory discrimination between two contexts can be caused by attenuation of the fear response in the conditioning context or by increasing the fear response in the neutral context. Our data indicate that activation of K3-GABA neurons does the latter; it decreases discrimination by driving fear generalization to a neutral context. Moreover, we find that K3-GABA activation does not interfere with the ability to acquire detailed fear memories. Instead, our results suggest that K3-GABA neurons play a specific role in the precision of memory recall. Because the activation of K3-GABA neurons severely decreases the net activity of CA3 neurons, our findings are consistent with the long-held notion that the CA3 is critically important for pattern completion during memory retrieval (Rolls and Treves 2024).

The hippocampus, and specifically the CA3, is particularly important for context-dependent memory precision (Olton, Walker et al. 1978, Kim and Fanselow 1992, Nakazawa, Quirk et al. 2002, Nakashiba, Young et al. 2008, Wiltgen, Zhou et al. 2010, Cherubini and Miles 2015, Guo, Soden et al. 2018). Moreover, previous work has detailed that as fear memories age, they lose their precision (Wiltgen and Silva 2007). Our results are consistent with a potential role for K3-GABA neurons in modulating the shift of memories from detailed to vague. In addition our results begin to fill a significant gap in understanding how nuanced control of CA3 circuits via different cellular inputs and outputs translates to contextual memory. Our work on K3-GABA neurons further supports the idea that CA3 circuits play a critical role in memory precision during recall and extends this knowledge by identifying a critical group of GABA neurons that may regulate this process by constraining CA3 neuron activity.

Interestingly, our results indicate that activating K3-GABA neurons sharply reduces CA3 activity but does not appreciably change either upstream DG or downstream CA1 activity as measured by cFos. This suggests that K3-GABA neurons specifically inhibit CA3-CA3 recurrent activity. Recurrent CA3 activity is thought to use an auto-associative network to enable recalling details and associations of contextual information (Leake, Zinn et al. 2021, Rolls and Treves 2024). But it has been difficult to experimentally determine the specific role of recurrent CA3 activity because manipulating CA3 itself affects all inputs and outputs simultaneously. PV neurons are known to synapse on or near the CA3 soma, which is thought to gate CA3 activity, but this would also be expected to affect all inputs and outputs simultaneously. In contrast, somatostatin GABA neurons are thought to synapse primarily on the distal CA3 dendrites that are innervated by the entorhinal cortex (Pelkey, Chittajallu et al. 2017, Honore, Khlaifia et al. 2021) and their activity may selectively modulate entorhinal inputs. Interestingly, the molecular identity of GABA neurons that specifically innervate middle dendritic regions that receive CA3 recurrent input is not specifically known but our structural and functional data suggest that K3-GABA neurons may fill this niche. Interestingly, CA3 neurons receive far more synapses from other CA3 neurons than any other cell type (Rolls and Treves 2024). If K3-GABA neurons specifically inhibit recurrent CA3-CA3 activity, this could explain why K3-GABA neurons more strongly inhibit CA3 as compared to PV neurons. More electrophysiological and anatomical work is needed to understand precisely how K3-GABA neurons are connected and how their intrinsic electrical properties compare to other GABA types, but changing K3-GABA neuron activity has potential to be a useful new avenue to fine-tune recurrent CA3-CA3 activity and thereby reveal its precise functions in learning and memory.

Kirrel3 is not a canonical marker for the study of GABA neurons, but it was shown that, in combination, synaptic CAMs are good predictors of functionally relevant groups of GABA neurons (Paul, Crow et al. 2017). Regardless of their classification, K3-GABA neurons comprise a significant proportion of GABA neurons in the hippocampus and K3-GABA neurons likely have clinical relevance because variations in the Kirrel3 gene, including copy number variations and missense mutations, are repeatedly associated with neurological disorders including autism, intellectual disability, and other mental illnesses (Bhalla, Luo et al. 2008, Kaminsky, Kaul et al. 2011, Ben-David and Shifman 2012, Guerin, Stavropoulos et al. 2012, Michaelson, Shi et al. 2012, Neale, Kou et al. 2012, Talkowski, Rosenfeld et al. 2012, De Rubeis, He et al. 2014, Iossifov, O’Roak et al. 2014, Wang, Guo et al. 2016, Li, Wang et al. 2017, Guo, Duyzend et al. 2019, Leblond, Cliquet et al. 2019, Taylor, Martin et al. 2020). Thus, patients with Kirrel3 gene variants likely have changes in circuits that express the Kirrel3 protein and rely on the function of Kirrel3. Since GABA neurons are well suited to fine-tune the net activity of neural circuits, understanding the role of Kirrel3-expressing GABA neurons on circuit function is relevant to developing potential therapeutic treatments for Kirrel3 disorders.

In sum, our work is one of the first to study a group of GABA neurons based on the expression of a molecule that drives synaptic connectivity. Cell type-specific studies of GABA neurons using cardinal markers like PV, somatostatin, and others enabled groundbreaking discoveries in circuit motifs, but it is becoming clear that even these tools label heterogeneous populations of GABA neurons with diverse transcriptomics and circuit functions (Yao, van Velthoven et al. 2021, Wu, Sevier et al. 2023, Chamberland, Grant et al. 2024). Our work demonstrates that K3-GABA neurons regulate memory precision, indicating that the generation of new tools to study distinct groups of neurons marked by a disease-linked molecule like Kirrel3 will propel our understanding of circuit function in healthy and disease states.

## Supporting information

https://docs.google.com/document/d/1jV392fM4zxgcAc-ALq1YfLW816VA8tpoTVeW0T1adK8/edit?usp=sharing

## Acknowledgments

We thank past and present members of the Williams lab for advice and technical help including Alyssa Johnson, Kara Graves and Randi Rawson. We thank Drs Lindsey Schwarz and Alex Hughes for sharing ConVERGD reagents prior to publication. This work was supported by the following funding sources: NIH T32HD007491 (A.T-O), NIH R25 NS130964 (C.M.P.), seed grants from the University of Utah (M.E.W.) and Brain Research Foundation (M.E.W.), the Thomas N. Parks Chair (M.E.W) and NIH grants U24 NS120055 (M.H.E), R01 MH125943 (M.E.W.) and R01 MH134515 (M.E.W. and J.G.H.).

## EXTENDED DATA FIGURE LEGENDS

**Extended Data Figure 3-1.** A) Examples of cues used for different contexts. B-D) Baseline metrics for K3-Flp;Gad2-Cre mice treated with either saline (gray) or DCZ (purple). n = 19 saline, 17 DCZ Each data point represents a mouse with males a triangle, females a circle. B) Fear acquisition curve for K3-Flp;Gad2-Cre mice shows the percent time spent freezing for 30 sec prior to each foot shock. Two-way ANOVA mixed effect test indicates a significant difference for shock number (indicating all mice conditioned normally) but no significant difference between treatment (saline vs DCZ). C) Baseline freezing equals the percent time freezing for each K3-Flp;Gad2-Cre mouse in the 3 minutes prior to the first foot shock. D) Baseline Activity Index shows overall motion of each K3-Flp;Gad2-Cre mouse in the 3 minutes prior to the first foot shock. E) Data from brain sections of Kirrel3-Flp; Gad2-Cre mice transfected with CV-hM3D virus. Plot indicates the percent area of hM3D-mCherry signal in CA3 versus outside CA3 as a measure of proper CA3 DREADD virus targeting and expression for each K3-Flp;Gad2-Cre mouse used in behavior experiments. Each row indicates a mouse, and each dot represents a brain section from that mouse. Error bars represent SEM. F-I) Data for WT WT mice treated with either saline (gray) or DCZ (cyan). n = 12 saline, 9 DCZ. Each data point represents a mouse with males a triangle, females a circle. F) shows % time freezing in each context, G) shows fear acquisition curves (two-way ANOVA mixed effect test indicates a significant difference for shock number indicating all mice conditioned normally but no significant difference between saline vs DCZ), H) shows the percent time spent freezing for 30 sec prior to each foot shock and I) shows the activity index for WT mice.

**Extended Data Figure 4-1.** A) Confocal images of adult triple heterozygous mice (K3-Flp;Gad2-Cre;RC-FLTG) stained for GFP to label K3-GABA neurons (green) and the indicated GABA neuron marker (magenta). B) Table showing the percent of K3-GABA neurons that express the indicated marker as assessed by immunostaining triple transgenic mice. n = 3 female adult mice. The total number of neurons counted for each marker is indicated. C) Representative image of a hippocampal section from an adult PV-Cre mouse expressing Cre-dependent hM3D-mCherry (magenta) and stained with anti-PV antibodies (green) indicates infected neurons express PV as expected. Arrowheads (yellow) point to overlap (white). D) Representative image of a PV-Cre mouse with a CA3 injection of the Cre-dependent DIO-hM3D-mCherry AAV. E) Fear acquisition curve shows freezing percent (%) for 30 sec prior to each foot shock. Two-way ANOVA mixed effect test indicates a significant difference for shock number indicating all mice conditioned normally but no significant difference between saline vs DCZ. F) Baseline freezing and G) activity index prior to foot shock for PV-Cre mice. For E-G, n = 19 saline, 16 DCZ. Error bars represent SEM. Each data point represents a mouse with males a triangle, females a circle.

**Extended Data Figure 5-1.** A) Fear acquisition curve shows freezing percent (%) for 30 sec prior to each foot shock for K3-GABA mice treated with saline or DCZ prior to the conditioning context only. Two-way ANOVA mixed effect test indicates a significant difference for shock number indicating all mice conditioned normally but no significant difference between saline vs DCZ. B) Baseline freezing and C) activity index prior to foot shock for animals in A. n = 15 saline, 10 DCZ. Error bars represent SEM, no significant differences as indicated. D) Fear acquisition curve shows freezing percent (%) for 30 sec prior to each foot shock for K3-GABA mice treated with saline or DCZ prior to Contexts A and C only on day 2. Two-way ANOVA mixed effect test indicates a significant difference for shock number indicating all mice conditioned normally but no significant difference between saline vs DCZ. E) Baseline freezing and F) activity index prior to foot shock for animals in D. n = 11 saline, 14 DCZ. Error bars represent SEM, no significant differences as indicated. In all cases, each data point represents a mouse with males a triangle, females a circle.

---

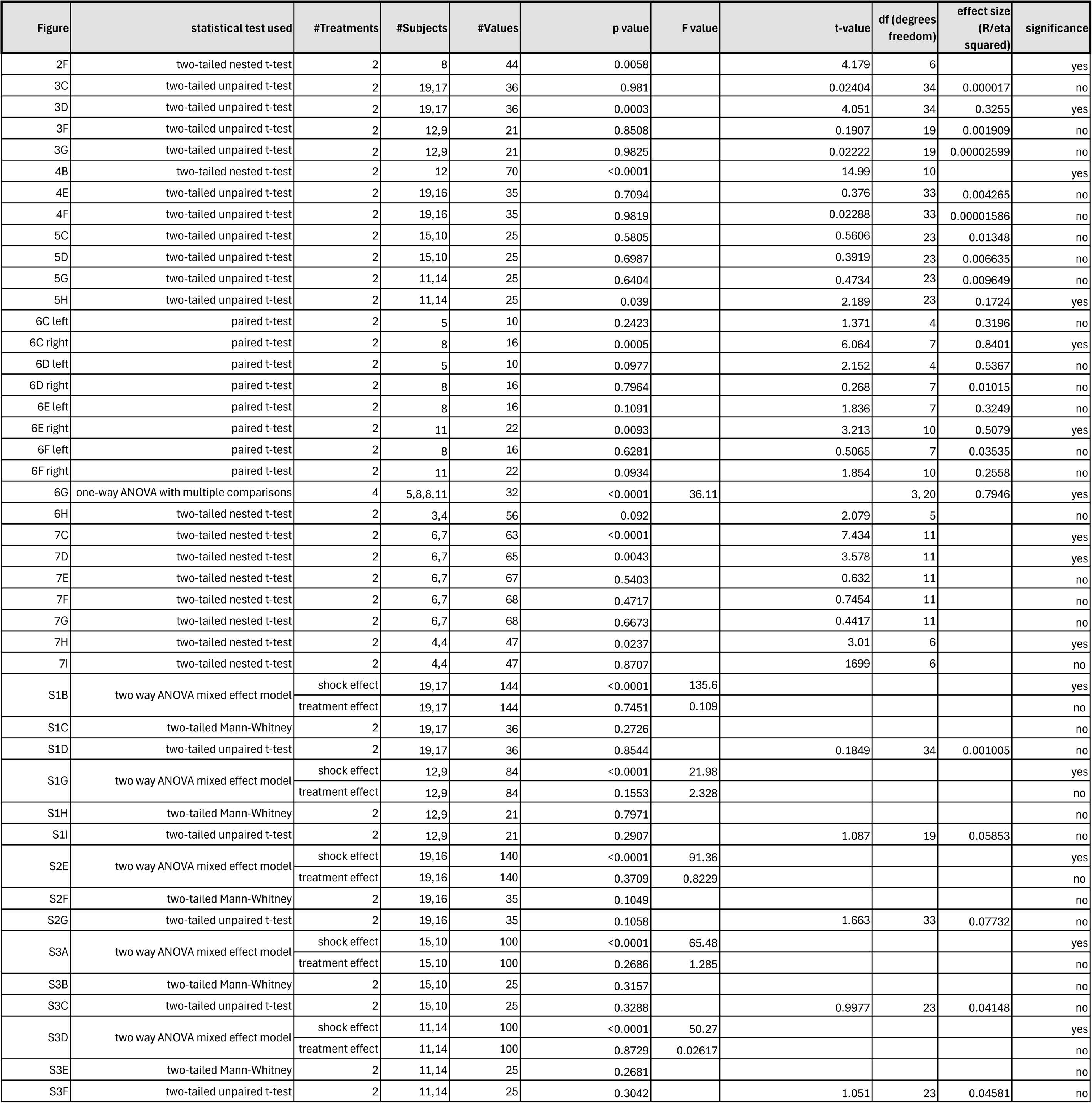

| Figure | statistical test used | #Treatments | #Subjects | #Values | p value | F value | t-value | df (degrees freedom) | effect size (R/eta squared) | significance |
| --- | --- | --- | --- | --- | --- | --- | --- | --- | --- | --- |
| 2F | two-tailed nested t-test | 2 | 8 | 44 | 0.0058 |  | 4.179 | 6 |  | yes |
| 3C | two-tailed unpaired t-test | 2 | 19,17 | 36 | 0.981 |  | 0.02404 | 34 | 0.000017 | no |
| 3D | two-tailed unpaired t-test | 2 | 19,17 | 36 | 0.0003 |  | 4.051 | 34 | 0.3255 | yes |
| 3F | two-tailed unpaired t-test | 2 | 12,9 | 21 | 0.8508 |  | 0.1907 | 19 | 0.001909 | no |
| 3G | two-tailed unpaired t-test | 2 | 12,9 | 21 | 0.9825 |  | 0.02222 | 19 | 0.00002599 | no |
| 4B | two-tailed nested t-test | 2 | 12 | 70 | <0.0001 |  | 14.99 | 10 |  | yes |
| 4E | two-tailed unpaired t-test | 2 | 19,16 | 35 | 0.7094 |  | 0.376 | 33 | 0.004265 | no |
| 4F | two-tailed unpaired t-test | 2 | 19,16 | 35 | 0.9819 |  | 0.02288 | 33 | 0.00001586 | no |
| 5C | two-tailed unpaired t-test | 2 | 15,10 | 25 | 0.5805 |  | 0.5606 | 23 | 0.01348 | no |
| 5D | two-tailed unpaired t-test | 2 | 15,10 | 25 | 0.6987 |  | 0.3919 | 23 | 0.006635 | no |
| 5G | two-tailed unpaired t-test | 2 | 11,14 | 25 | 0.6404 |  | 0.4734 | 23 | 0.009649 | no |
| 5H | two-tailed unpaired t-test | 2 | 11,14 | 25 | 0.039 |  | 2.189 | 23 | 0.1724 | yes |
| 6C left | paired t-test | 2 | 5 | 10 | 0.2423 |  | 1.371 | 4 | 0.3196 | no |
| 6C right | paired t-test | 2 | 8 | 16 | 0.0005 |  | 6.064 | 7 | 0.8401 | yes |
| 6D left | paired t-test | 2 | 5 | 10 | 0.0977 |  | 2.152 | 4 | 0.5367 | no |
| 6D right | paired t-test | 2 | 8 | 16 | 0.7964 |  | 0.268 | 7 | 0.01015 | no |
| 6E left | paired t-test | 2 | 8 | 16 | 0.1091 |  | 1.836 | 7 | 0.3249 | no |
| 6E right | paired t-test | 2 | 11 | 22 | 0.0093 |  | 3.213 | 10 | 0.5079 | yes |
| 6F left | paired t-test | 2 | 8 | 16 | 0.6281 |  | 0.5065 | 7 | 0.03535 | no |
| 6F right | paired t-test | 2 | 11 | 22 | 0.0934 |  | 1.854 | 10 | 0.2558 | no |
| 6G | one-way ANOVA with multiple comparisons | 4 | 5,8,8,11 | 32 | <0.0001 | 36.11 |  | 3, 20 | 0.7946 | yes |
| 6H | two-tailed nested t-test | 2 | 3,4 | 56 | 0.092 |  | 2.079 | 5 |  | no |
| 7C | two-tailed nested t-test | 2 | 6,7 | 63 | <0.0001 |  | 7.434 | 11 |  | yes |
| 7D | two-tailed nested t-test | 2 | 6,7 | 65 | 0.0043 |  | 3.578 | 11 |  | yes |
| 7E | two-tailed nested t-test | 2 | 6,7 | 67 | 0.5403 |  | 0.632 | 11 |  | no |
| 7F | two-tailed nested t-test | 2 | 6,7 | 68 | 0.4717 |  | 0.7454 | 11 |  | no |
| 7G | two-tailed nested t-test | 2 | 6,7 | 68 | 0.6673 |  | 0.4417 | 11 |  | no |
| 7H | two-tailed nested t-test | 2 | 4,4 | 47 | 0.0237 |  | 3.01 | 6 |  | yes |
| 7I | two-tailed nested t-test | 2 | 4,4 | 47 | 0.8707 |  | 1699 | 6 |  | no |
| S1B | two way ANOVA mixed effect model | shock effect | 19,17 | 144 | <0.0001 | 135.6 |  |  |  | yes |
|  |  | treatment effect | 19,17 | 144 | 0.7451 | 0.109 |  |  |  | no |
| S1C | two-tailed Mann-Whitney | 2 | 19,17 | 36 | 0.2726 |  |  |  |  | no |
| S1D | two-tailed unpaired t-test | 2 | 19,17 | 36 | 0.8544 |  | 0.1849 | 34 | 0.001005 | no |
| S1G | two way ANOVA mixed effect model | shock effect | 12,9 | 84 | <0.0001 | 21.98 |  |  |  | yes |
|  |  | treatment effect | 12,9 | 84 | 0.1553 | 2.328 |  |  |  | no |
| S1H | two-tailed Mann-Whitney | 2 | 12,9 | 21 | 0.7971 |  |  |  |  | no |
| S1I | two-tailed unpaired t-test | 2 | 12,9 | 21 | 0.2907 |  | 1.087 | 19 | 0.05853 | no |
| S2E | two way ANOVA mixed effect model | shock effect | 19,16 | 140 | <0.0001 | 91.36 |  |  |  | yes |
|  |  | treatment effect | 19,16 | 140 | 0.3709 | 0.8229 |  |  |  | no |
| S2F | two-tailed Mann-Whitney | 2 | 19,16 | 35 | 0.1049 |  |  |  |  | no |
| S2G | two-tailed unpaired t-test | 2 | 19,16 | 35 | 0.1058 |  | 1.663 | 33 | 0.07732 | no |
| S3A | two way ANOVA mixed effect model | shock effect | 15,10 | 100 | <0.0001 | 65.48 |  |  |  | yes |
|  |  | treatment effect | 15,10 | 100 | 0.2686 | 1.285 |  |  |  | no |
| S3B | two-tailed Mann-Whitney | 2 | 15,10 | 25 | 0.3157 |  |  |  |  | no |
| S3C | two-tailed unpaired t-test | 2 | 15,10 | 25 | 0.3288 |  | 0.9977 | 23 | 0.04148 | no |
| S3D | two way ANOVA mixed effect model | shock effect | 11,14 | 100 | <0.0001 | 50.27 |  |  |  | yes |
|  |  | treatment effect | 11,14 | 100 | 0.8729 | 0.02617 |  |  |  | no |
| S3E | two-tailed Mann-Whitney | 2 | 11,14 | 25 | 0.2681 |  |  |  |  | no |
| S3F | two-tailed unpaired t-test | 2 | 11,14 | 25 | 0.3042 |  | 1.051 | 23 | 0.04581 | no |

| Reagent | RRID number |
| --- | --- |
| pAAV-hSyn-DIO-hM3D(Gq)-mCherry | Addgene_44361 |
| PV-Cre mouse (Strain #:017320 Jackson Laboratory) | IMSR_JAX:017320 |
| Olympus VS200 Slide Scanner | SCR_024783 |
| MATLAB | SCR_001622 |
| KiloSort | SCR_016422 |
| Python Programming Language | SCR_008394 |
| C57BL/6J wildtype mice (JAX stock #000664) | RRID:IMSR_JAX:000664 |
| Gad2-IRES-Cre (JAX stock #010802) | RRID:IMSR_JAX:010802 |
| RC::FLTG (JAX stock #026932) | RRID:IMSR_JAX:026932 |
| goat anti-GFP 1:5000 (Abcam, ab6673) | RRID:AB_305643 |
| rabbit anti-cFos 1:500 (Santa Cruz Biotech, sc-253) | RRID:AB_2231996 |
| rabbit anti-cFos 1:500 (Synaptic Systems, 226008) | RRID:AB_2891278 |
| rabbit anti-calretinin 1:2000 (Swant, 7699/3) | RRID:AB_10000321 |
| mouse anti-parvalbumin 1:5000 (Swant, PV235) | RRID:AB_3698492 |
| rat anti-somatostatin 1:500 (Chemicon, MAB354) | RRID:AB_2255365 |
| rabbit anti-calbindin d28k 1:2000 (Swant, CB38a) | RRID:AB_3107026 |
| rabbit anti-VIP 1:500 (Immunostar) | RRID:AB_572270 |
| rabbit anti-PV 1:250 (Swant, PV27a) | RRID:AB_2631173 |
| rabbit anti-RFP 1:1000 (Fisher, NBP267102) | RRID:AB_560939 |
| goat anti-mCherry (Novus, NBP3-05558) | RRID:AB_3534099 |
| (AAV8)CV.hSyn.hM3D.HA | RRID:Addgene_218758 |
| (AAV9)hSyn.DIO.hM3D.mCherry (Addgene 44361) | RRID:Addgene_44361 |
| hSyn.Cre-on/Flp-on.YFP (Addgene 55650) | RRID:Addgene_55650 |
| Zeiss 980 | RRID:SCR_025048 |
| FIJI | RRID:SCR_002285 |
| DRAQ5 (1:1000, Cell Signaling Technology) | RRID:AB_2869620 |

