## Supplementary material for "Inhibitory neurons marked by the connectivity molecule Kirrel3 regulate memory precision": https://docs.google.com/document/d/1jV392fM4zxgcAc-ALq1YfLW816VA8tpoTVeW0T1adK8/edit?usp=sharing

### Extended Data Figure 3-1

**A**

| | Context A | Context B | Context C |
|---|---|---|---|
| Floor | Even bars | Flat insert | Uneven Bars |
| Roof | Standard | Standard | Triangular insert |
| Scent | SimpleGreen | Simple Green | Windex |
| Sound | Fan | Fan | None |
| Light | Yes | Yes | No light |
| Transport | Bag | Covered Cart | Uncovered Cart |

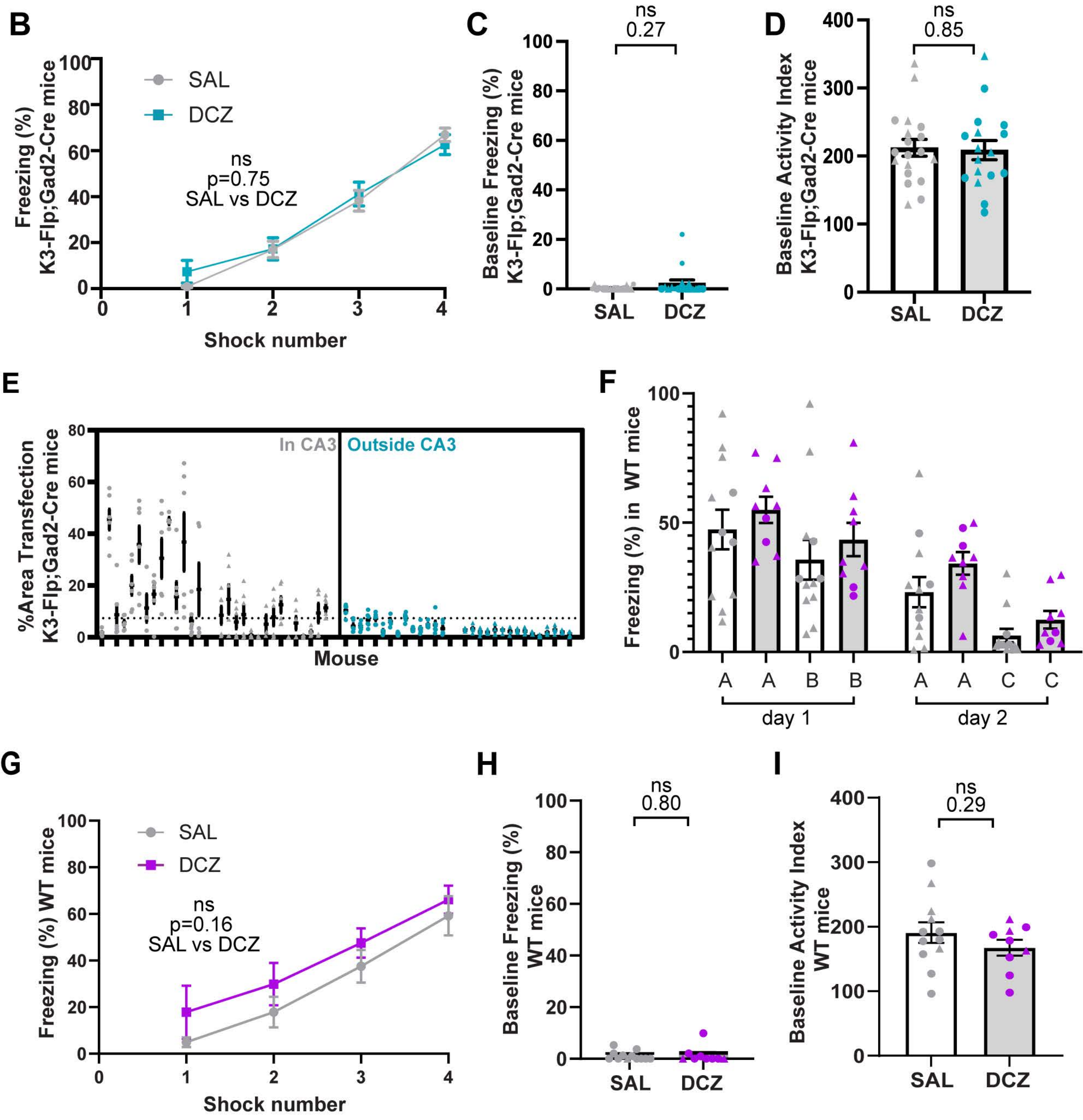

### Extended Data Figure 4-1

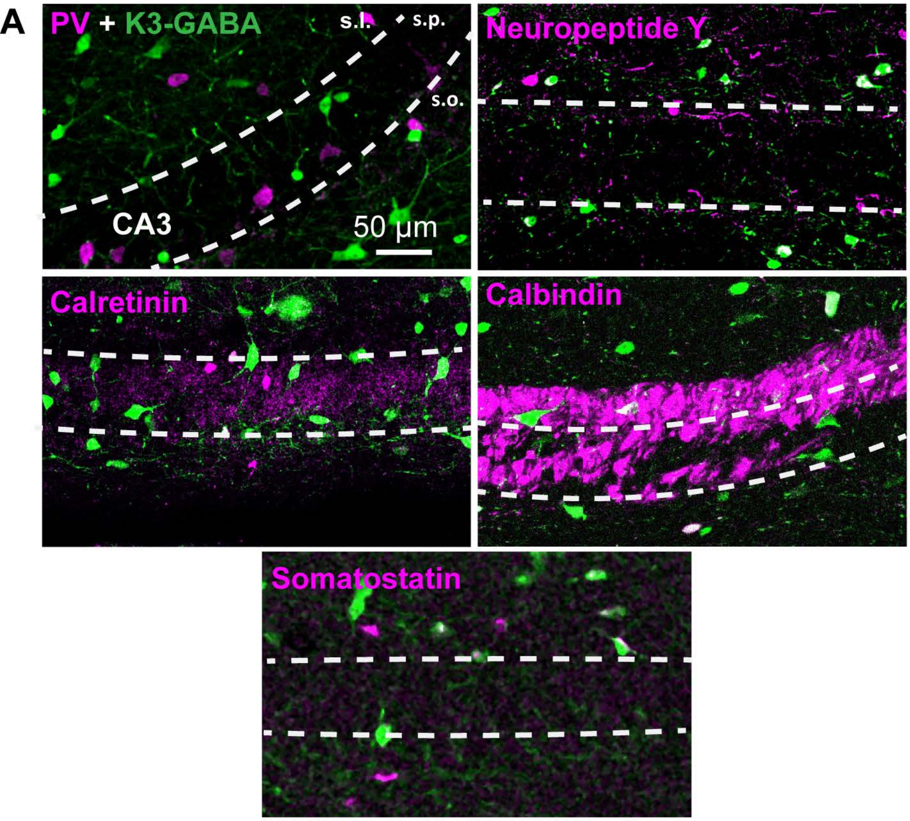


B

| GABA Marker | %K3-GABA |
|---|---|
| Calretinin | 7% (179/2680) |
| Somatostatin | 19% (298/1573) |
| Neuropeptide Y | 32% (656/2024) |
| Parvalbumin | 9% (196/2124) |
| Calbindin | 23% (89/292) |

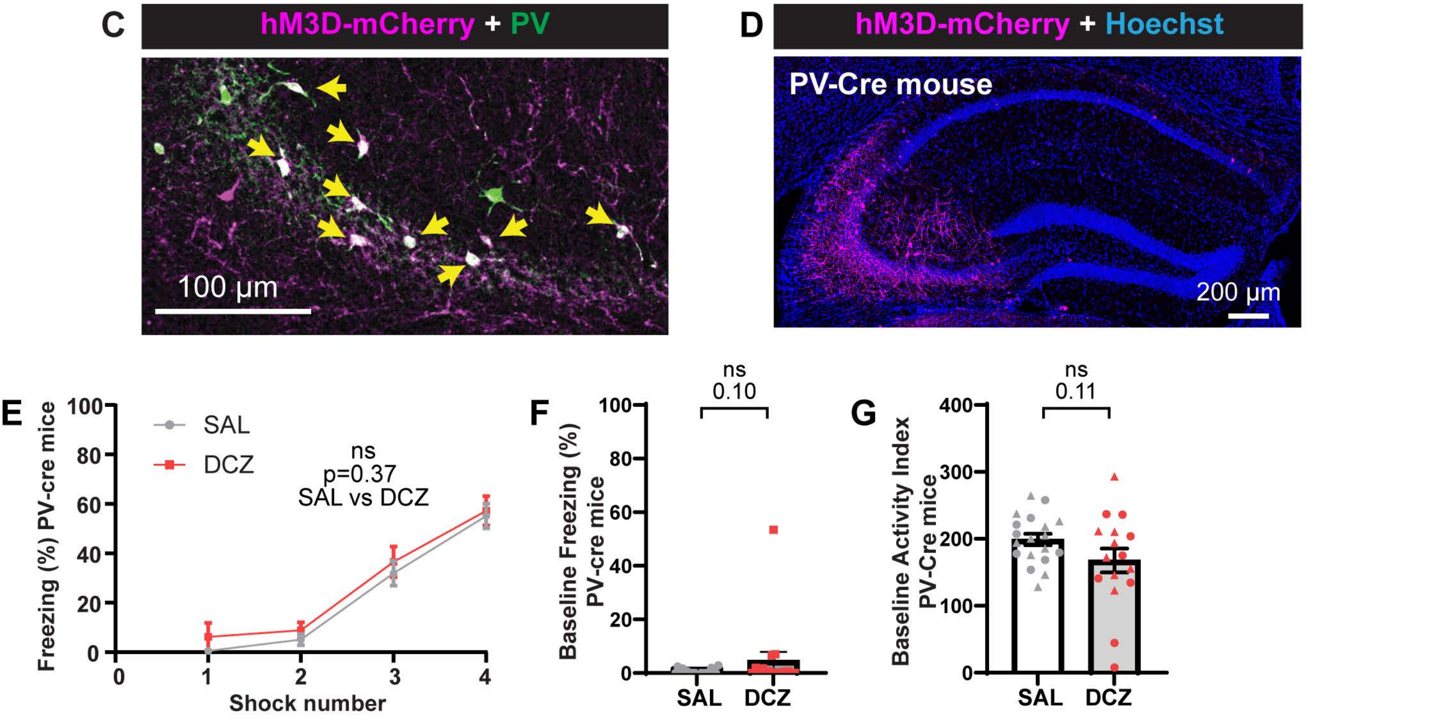

### Extended Data Figure 5-1

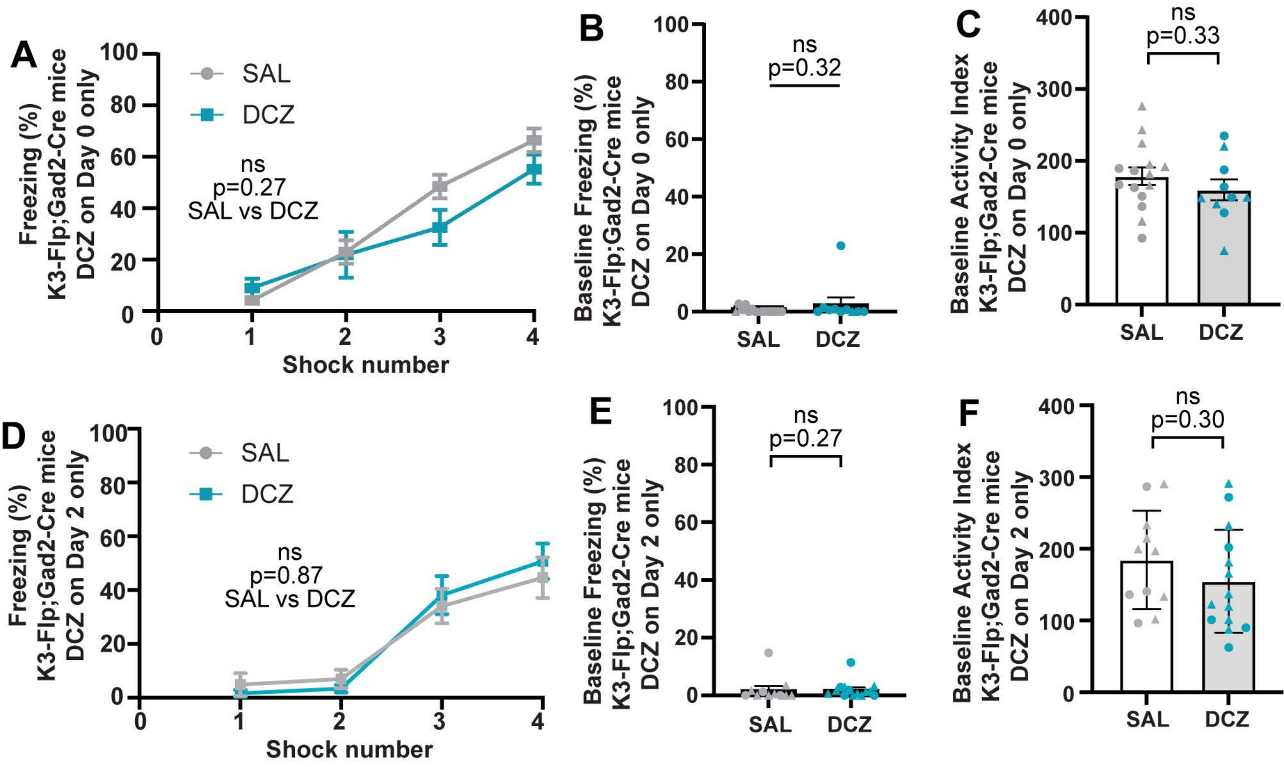
